# Breed-Specific Characteristics of Beef Are Partly Retained in Cultivated Meat

**DOI:** 10.1101/2025.07.24.666483

**Authors:** Chika Nakadozono, Fiona Louis, Asuka Yamada, Miki Mizukami, Yoshihiro Kodama, Shiro Kitano, Takuya Sugiura, Akiko Takasuga, Koji Nakade, Michiya Matsusaki

## Abstract

Cultivated meat (CM) is being developed as an alternative to conventional meat, but whether characteristics associated with the source breed are reflected in CM remains unclear. Here, we systematically compared the biological and physicochemical properties of Wagyu, Wagyu–Holstein crossbred (Crossbreed), and Holstein beef with those of cultivated muscle and fat fibers fabricated by three-dimensional (3D) bioprinting using bovine satellite cells (bSCs) and bovine adipose-derived stem cells (bADSCs) isolated from the respective cattle. Sensory evaluation was conducted on conventional beef samples only. The cultivated fibers exhibited breed-associated differences in amino acid and fatty acid profiles, aroma-related compounds, and gene expression. In particular, cells derived from Wagyu showed distinct expression profiles related to adipogenesis and lipid metabolism, whereas those derived from Holstein showed distinct profiles related to myogenesis and extracellular matrix. However, the cultivated fibers did not fully reproduce the composition of native beef tissues. These findings indicate that some breed-associated characteristics are reflected in cultivated fibers and that cell origin is an important factor in determining CM properties. This study provides a basis for designing CM with selected breed-associated characteristics.

**TOC:** 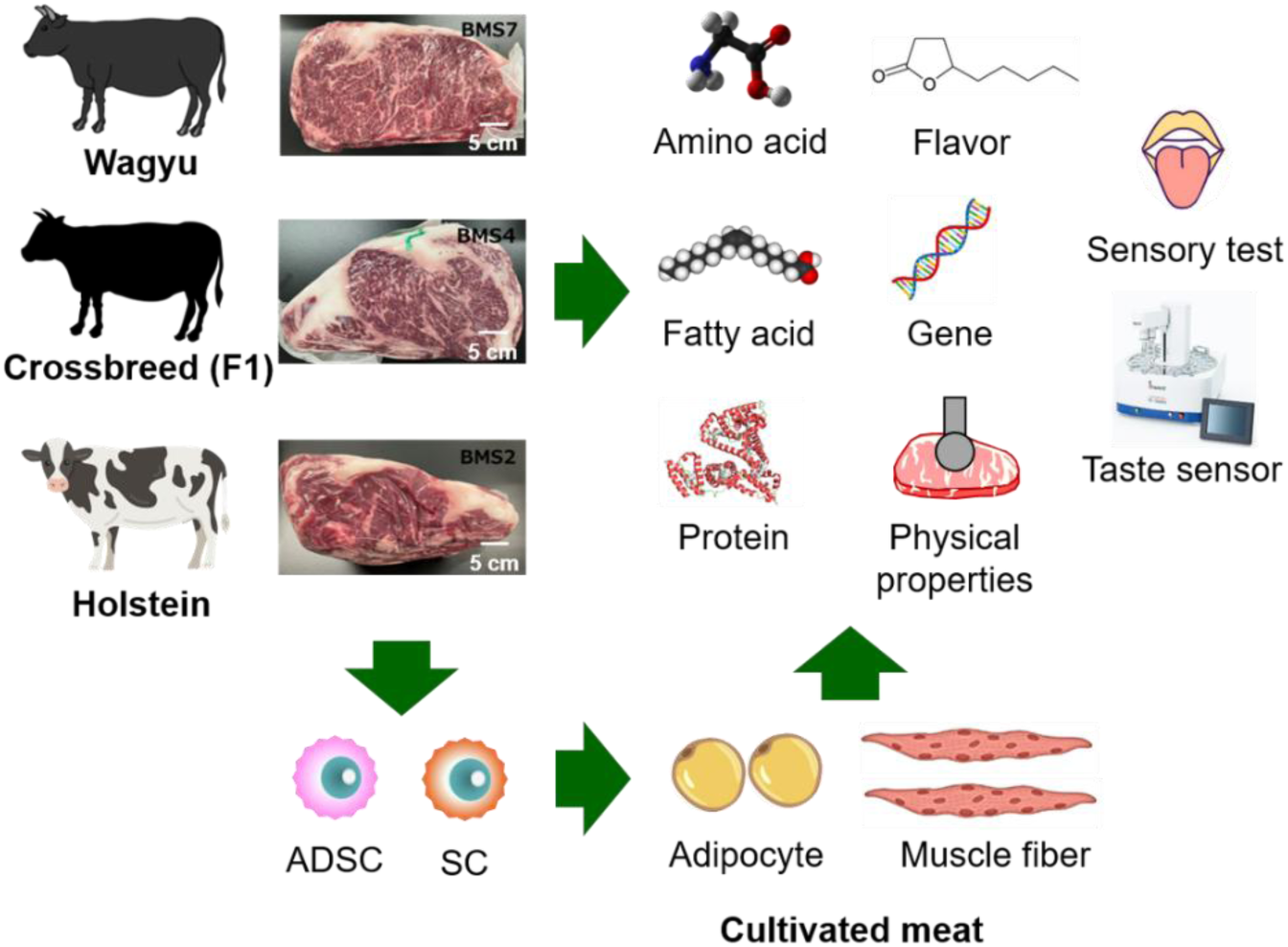

## Introduction

Over the past decade, cultivated meat (CM) has attracted considerable attention from the perspectives of ethics, economics, environmental sustainability, and public health, although several challenges remain to be addressed. The Singapore Food Agency defines CM as “meat developed from animal cell culture, where the production process involves growing the selected cell lines (or stem cells) in a bioreactor. These cells are grown in a suitable growth medium, and subsequently onto a scaffold to produce products resembling meat muscle”^1^. Commercial products containing plant-based proteins and lysates of chicken fibroblast or myoblast cells have recently emerged. Unlike conventional meat alternatives, CM has the potential to reproduce the characteristics of conventional meat by controlling its cellular composition and the resulting taste, flavor, and texture.

Various bio-manufacturing methods have been developed for CM production. Scaffolds with cell-adhesive properties play an important role in supporting the survival and expansion of adherent satellite cells (SCs), which can subsequently differentiate into myoblasts and myotubes. In vitro replication of cell–extracellular matrix (ECM) interactions has been achieved using scaffolding materials with inherent integrin-binding sites, including collagen^2^, gelatin^3^, and fibrin^4^. Biologically inert materials, such as polysaccharides (e.g., alginate, gellan gum, and agar) and plant-based proteins (e.g., soy and pea proteins), generally require functionalization with integrin ligands, such as arginine-glycine-aspartate (RGD) motifs^5^ or peptides derived from laminin and fibronectin^6^. Because one of the objectives of CM production is to reduce animal slaughter and the use of animal-derived products, the source and production methods of scaffolding materials are also important considerations.

Various tissue engineering approaches, including top-down scaffold-based methods, bottom-up cell-sheet stacking, and three-dimensional (3D) bioprinting, have been investigated for CM production, as summarized in recent reviews^7,8^. Top-down approaches need a scaffold including microcarriers^9^, fibrous spun matrix^10^, and decellularized scaffolds from plants or vegetables^11^ for the growth and differentiation of cells. Bottom-up cell-sheet stacking enables the construction of CM with high cell density through a scaffold-free approach^12^. In contrast, 3D bioprinting, particularly extrusion-based bioprinting, enables the fabrication of meat-like structures with controlled architecture and composition. By directly combining cells and scaffold materials during printing, their densities, compositions, and spatial distributions can be controlled. To construct CM with tissue-like structures, high cell density (>10⁷ cells/mL) and unidirectionally aligned, mature myotubes are desirable. However, the strong contractile forces generated by differentiated muscle cells can deform the scaffolds. To address this issue, we previously developed an anchoring technology using tendon-like, collagen-based tough gels for 3D-printed bovine satellite cell (bSC) and adipose-derived stem cell (bADSC) fibers^4^. The resulting muscle and fat fibers could be assembled into marbled structures resembling Wagyu beef, with the muscle-to-fat ratio being readily controlled.

Evaluating the organoleptic properties and nutritional composition of CM is essential for determining how closely its characteristics resemble those of conventional meat^13^. Previous studies have reported that CM generally exhibits higher water content and softer mechanical properties than conventional meat^9,14^, often accompanied by lower protein and amino acid contents^12,15^. However, direct comparisons among reported CM studies remain challenging because of differences in cell sources (e.g., chicken, cattle, and fish), scaffold materials (e.g., animal-derived ECM, decellularized plant materials, and synthetic polymers), culture and differentiation media, culture periods, and analytical methods. In particular, it remains unclear to what extent characteristics associated with the source breed are reflected in CM when cells derived from different cattle breeds are processed using the same fabrication method. A systematic comparison under controlled fabrication conditions is therefore needed to understand the relationship between the characteristics of conventional beef and those of CM produced from cells derived from the corresponding cattle.

In this study, we systematically investigated the physicochemical and biological characteristics of Wagyu (W), Wagyu–Holstein crossbred (Crossbreed, C), and Holstein (H) beef, and compared them with those of muscle and fat fibers fabricated by 3D bioprinting using bSCs and bADSCs isolated from the respective cattle. We evaluated nutritional composition, fatty acid profiles, aroma-related compounds, transcriptomic and proteomic profiles, and physical properties of the conventional beef and cultivated fibers. Sensory evaluation was conducted on conventional beef samples to characterize breed-associated differences in sensory properties. By comparing the properties of cultivated fibers produced using the same fabrication method, we aimed to determine whether and to what extent breed-associated characteristics are reflected in CM (Fig. 1).

**Fig. 1.**
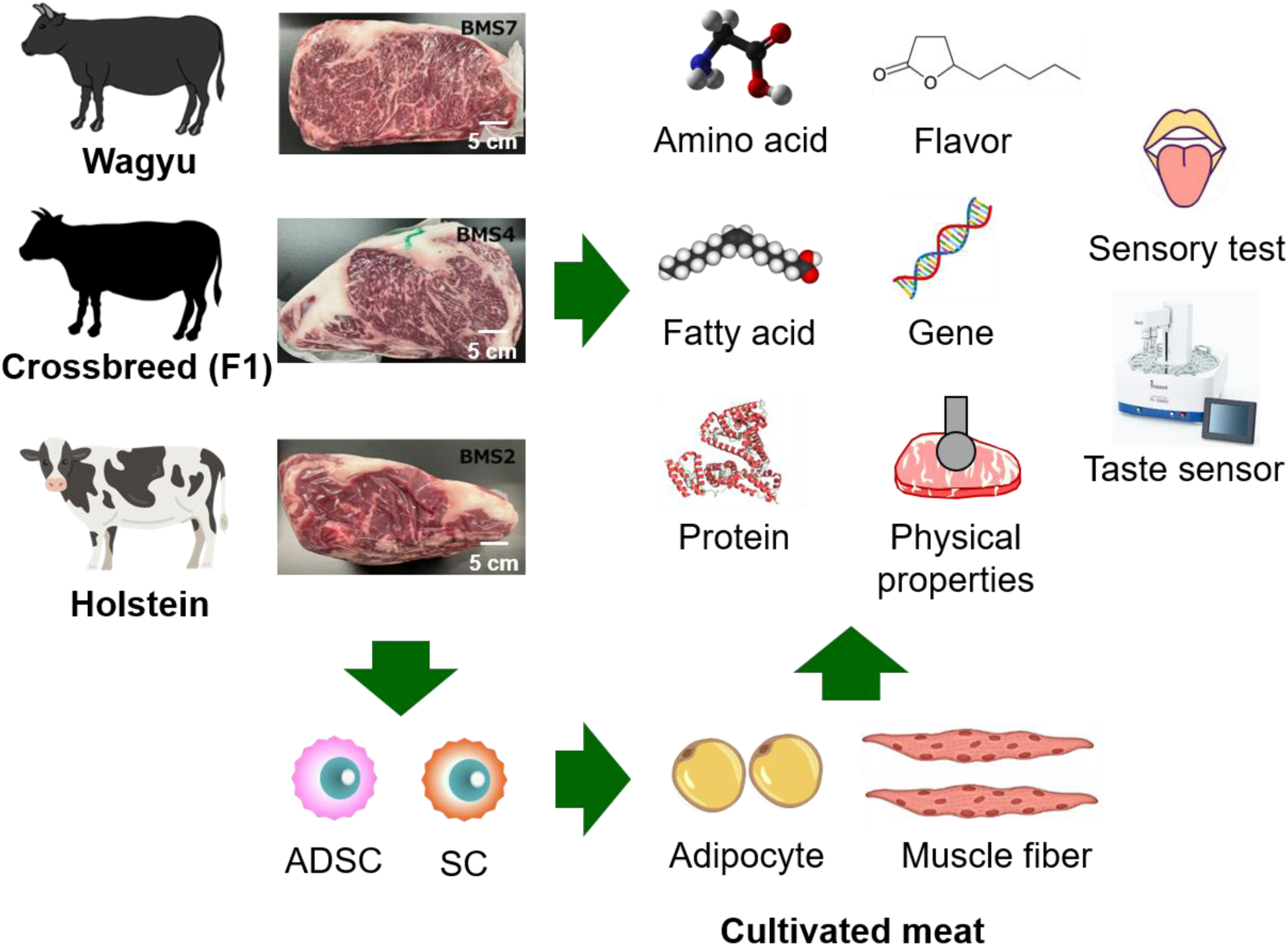
Schematic illustration of this study. Biological and physicochemical analyses of the meats of Wagyu (W), Crossbreed (C) and Holstein (H) were performed to understand the different characteristics. Adipose derived stem cells (ADSCs) and satellite cells (SCs) were isolated from each fresh meat and then differentiated into adipocyte and muscle fibers after the preculture. The same biological and physicochemical analyses were performed on the obtained adipocytes and muscle fibers to understand how the differentiated tissues differ from the original meats. Sensory evaluation was performed only on conventional beef samples, not on cultivated meat samples.

## Results

### Physical properties and nutrient composition of conventional beef

The physical properties of Wagyu, Crossbreed, and Holstein beef were evaluated by texture profile analysis, including hardness, chewiness, cohesiveness, and springiness, in both raw and boiled samples (Fig. 2a,b). Boiling increased all texture parameters except springiness. In particular, boiled Holstein beef was significantly harder than its raw counterpart (3.27-fold), whereas the corresponding increases were 1.29-fold and 2.57-fold for Crossbreed and Wagyu, respectively. Histological images obtained by hematoxylin and eosin (HE) staining suggested that Wagyu and Crossbreed beef had a more homogeneous distribution of adipose tissue than Holstein beef. In addition, Holstein beef showed a greater percentage of weight loss after boiling than Wagyu and Crossbreed beef (Supplementary Fig. S1a,b).

**Fig. 2.**
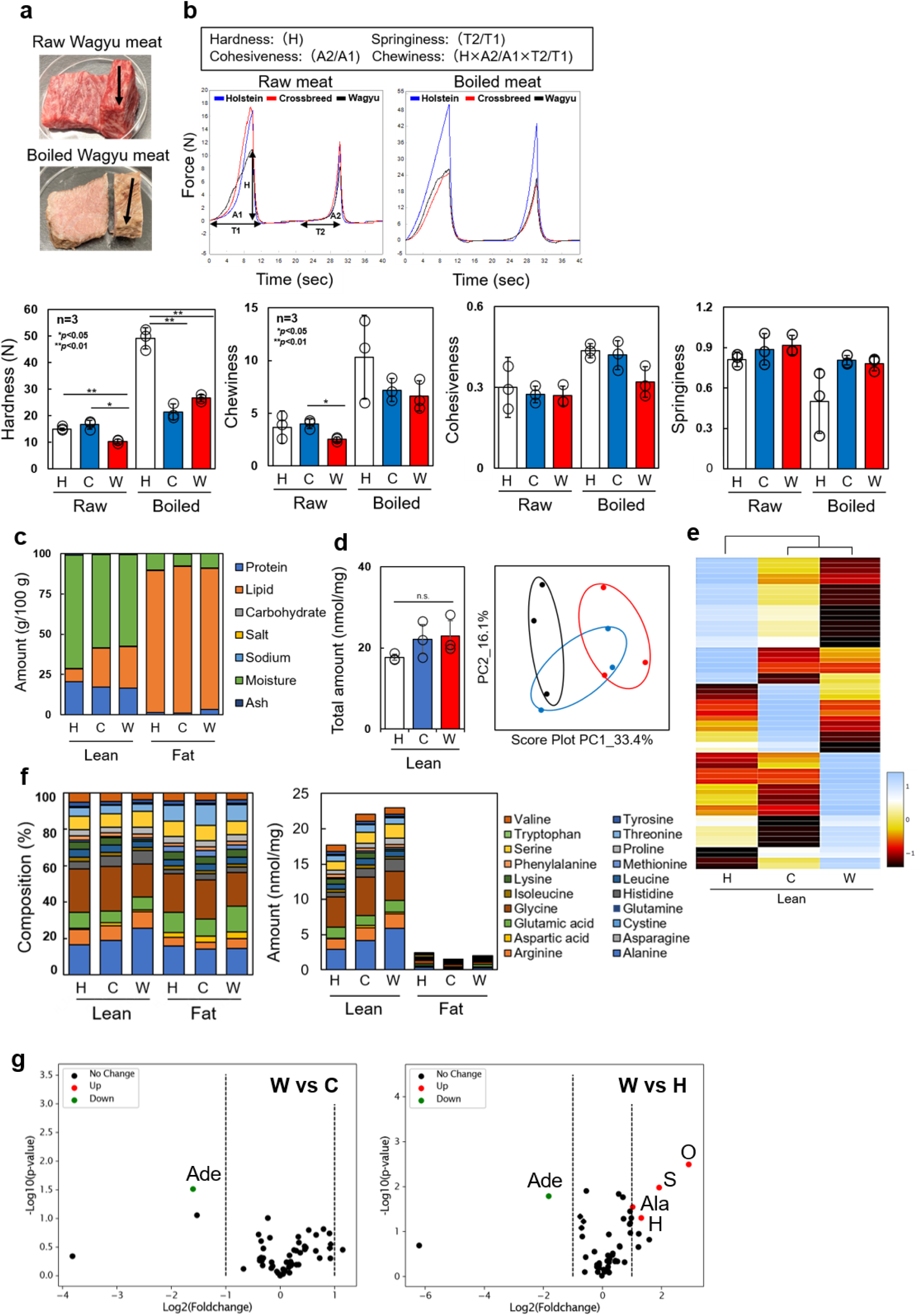
Physical properties and nutrient composition of each meat. a, Photographs of raw and boiled Wagyu meat. Black arrows indicate muscle fiber direction, and compression experiments were performed vertical direction of the arrows. b, Texture profile analysis of raw and boiled meats (left) and the data of each parameter (right) (n=3). c, Mean nutrient value of lean and fat of each beef meat, respectively (n=3). d, Total amount and principal component analysis (PCA) of amino acids and nucleic acid-related substances of the lean meats (n=3). Black is Holstein, blue is Crossbreed and red is Wagyu, respectively. e, Heat map analysis of free amino acid of each beef meat. f, Composition and concentration of free amino acid of 100g of mixed of lean and fat of each beef meat, respectively (n=3). g, Volcano plots of free an amino acid, nucleoside and vitamin in lean of W vs C and W vs H. Ade: Adenosine monophosphate, O: Ornithine, S: Symmetric dimethylarginine, Ala: Alanine, H: Histidine, respectively.

To investigate differences in nutritional composition, the nutrient contents and free amino acid profiles of the lean and fat portions of each beef type were analyzed (Fig. 2c,d). Wagyu beef had the highest lipid content in the lean portion (25.75 g/100 g), which was 3.23-fold higher than that of Holstein beef, whereas Holstein beef had the highest protein content (20.35 g/100 g). In contrast, the lipid contents of the fat portions were similar among the three beef types, ranging from 87.6 to 91.0 g/100 g. Although Wagyu beef showed the highest total free amino acid content (73.4 nmol/mg), the difference was not statistically significant. Principal component analysis (PCA) of free amino acids and nucleic acid-related compounds indicated differences between Wagyu and Holstein beef, with Crossbreed samples showing an intermediate distribution (Fig. 2d). Heat map analysis of free amino acids in the lean portions also revealed different profiles among the three beef types, with Wagyu and Crossbreed showing greater similarity to each other than to Holstein (Fig. 2e).

The free amino acid composition of samples prepared by mixing the lean and fat portions of each beef type was also evaluated (Fig. 2f). Alanine and glycine, both of which contribute to sweetness, were among the predominant amino acids in the lean and fat portions of all three beef types. Volcano plot analysis of free amino acids, nucleosides, and vitamins in the lean portions revealed that adenosine monophosphate was less abundant in Wagyu than in Crossbreed and Holstein, whereas ornithine, symmetric dimethylarginine, alanine, and histidine were more abundant in Wagyu (Fig. 2g). These results demonstrate breed-associated differences in the nutritional composition and free amino acid profiles of conventional beef.

### Fatty acid and aroma profiles of conventional beef

Because fatty acid composition and aroma compounds contribute to the flavor characteristics of beef, the total fatty acid contents and the proportions of saturated fatty acids (SFAs), monounsaturated fatty acids (MUFAs), and polyunsaturated fatty acids (PUFAs) were analyzed by gas chromatography–mass spectrometry (GC– MS) (Fig. 3a,b). Wagyu beef showed the highest total fatty acid content, particularly in the SFA and MUFA fractions, whereas no corresponding increase was observed in the PUFA fraction. Analysis of individual fatty acids showed that oleic acid (C18:1n9c) was the predominant fatty acid in all three beef types, followed by palmitic acid (C16:0) and stearic acid (C18:0) (Fig. 3c). The proportion of oleic acid was highest in Wagyu beef (44%), compared with approximately 40% in Crossbreed and Holstein beef. The proportions of palmitic and stearic acids were similar among the three beef types, at approximately 25–30% and 15–20%, respectively.

**Fig. 3.**
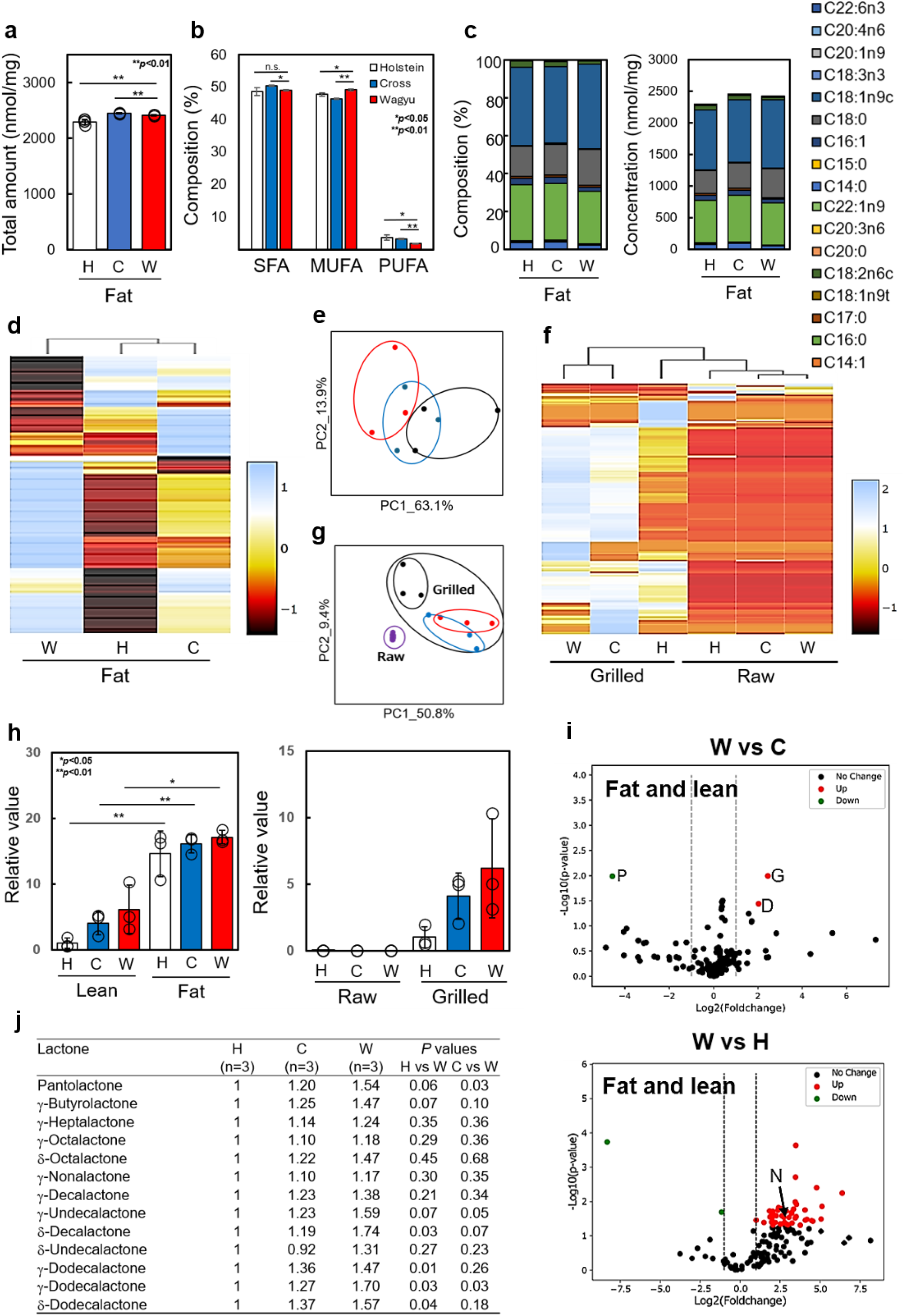
Fatty acid and aroma analyses of each meat. **a**, Total amount of fatty acids of each beef meat (n=3). **b**, Composition of saturated fatty acids (SFA), monounsaturated fatty acids (MUFA) and polyunsaturated fatty acids (PUFA) of each beef meat (n=3). **c**, Composition and concentration of fatty acids of fat in each beef meat, respectively (n=3). A heat map (**d**) and PCA analysis (**e**) of aroma components of fat tissues of each meat. Black is Holstein, blue is Crossbreed and red is Wagyu in **e**, respectively. A heat map (**f**) and PCA analysis (**g**) of aroma components of fat tissues in grilled and raw each meat. **h**, The relative secreted value of γ-nonalactone from the lean and fat tissues of each beef meat (left), and the comparison of it with lean and fat tissues w/o or w/ grilling (right), respectively (n=3). **i**, Volcano plots of secreted aroma components of grilled lean and fat tissues in between W vs. C (left) and W vs. H (right), respectively. **j**, Summary of the relative values of secreted lactone species from the grilled fat tissues in Wagyu and Crossbreed meets as compared to those from Holstein.

The aroma profiles of the fat portions were further evaluated by heat map and PCA analyses (Fig. 3d,e), and the detected aroma compounds are summarized in Supplementary Fig. S2. Crossbreed and Holstein showed similar aroma profiles, whereas Wagyu exhibited a distinct profile. PCA also indicated a difference between Wagyu and Holstein, with Crossbreed showing an intermediate distribution, consistent with the trends observed in the analyses of amino acids and nucleic acid-related compounds (Fig. 2d). Comparison of raw and grilled samples revealed that grilling increased the release of aroma compounds, particularly in Wagyu and Crossbreed beef (Fig. 3f). PCA showed a wider distribution of the grilled samples than of the raw samples. The distributions of grilled Wagyu and Crossbreed samples largely overlapped, whereas Holstein samples were separated from them (Fig. 3g).

To investigate characteristic aroma compounds in Wagyu beef, we focused on γ-nonalactone, which has been associated with the sweet aroma of Wagyu beef^16^. The relative amount of γ-nonalactone detected from the fat portions was higher than that from the lean portions, and Wagyu showed the highest values in both portions among the three beef types (Fig. 3h, left). Grilling increased the detected amount of γ-nonalactone in both lean and fat portions, with a particularly pronounced increase in Wagyu beef (6.16-fold) (Fig. 3h, right). Volcano plot analysis of aroma compounds released from grilled lean and fat portions revealed differences between Wagyu and Crossbreed, as well as between Wagyu and Holstein. Lauric acid was more abundant in Wagyu in both comparisons, while 2-heptanone was higher and homofureanol and (2E,6Z)-nona-2,6-dienal were lower in Wagyu than in Crossbreed (Fig. 3i). The relative amounts of lactone species released from grilled fat portions are summarized in Fig. 3j. Wagyu showed the highest values for all the lactones examined, particularly γ-undecalactone (1.59-fold), δ-decalactone (1.74-fold), and γ-dodecalactone (1.70-fold), compared with Holstein. These results indicate that Wagyu beef has distinct fatty acid and aroma profiles, including higher levels of several lactone compounds associated with sweet aroma.

### Transcriptomic profiles of bovine cells and sensory evaluation of conventional beef

The gene expression profiles of bovine satellite cells (bSCs) and bovine adipose-derived stem cells (bADSCs) isolated from the three cattle types were analyzed by RNA sequencing (RNA-seq) (Fig. 4). PCA showed that the expression profiles of Wagyu- and Crossbreed-derived cells were relatively similar, with some overlap, whereas those of Holstein-derived cells were more distinct (Fig. 4a). Volcano plots comparing Wagyu with Crossbreed or Holstein are shown in Fig. 4b. In bSCs, the numbers of significantly upregulated and downregulated genes in Crossbreed relative to Wagyu were 338 and 327, respectively. The corresponding numbers in Holstein relative to Wagyu were higher, at 803 and 1,393, respectively. In bADSCs, the numbers of upregulated and downregulated genes were 1,345 and 999 for Crossbreed relative to Wagyu, and 1,302 and 1,186 for Holstein relative to Wagyu.

**Fig. 4.**
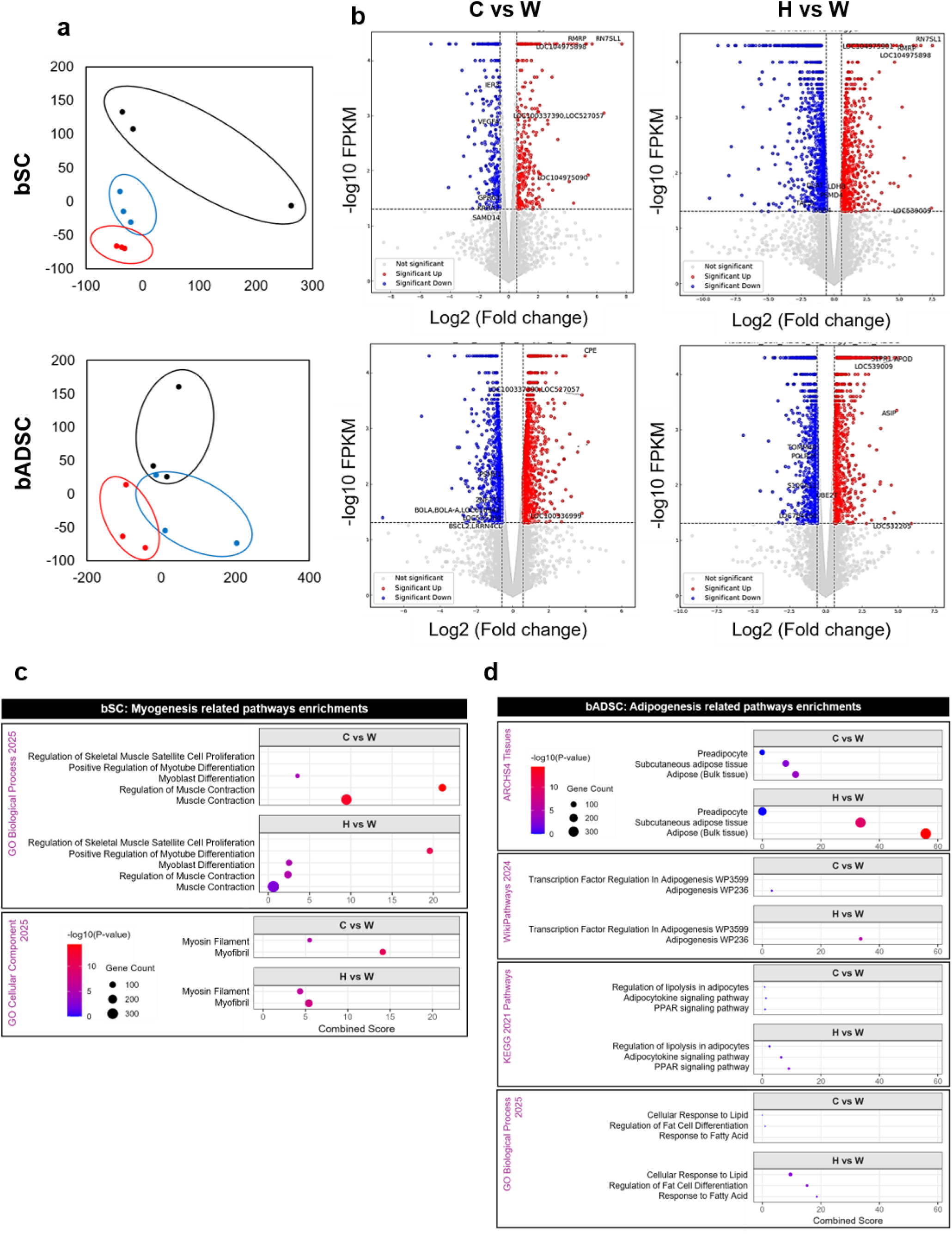
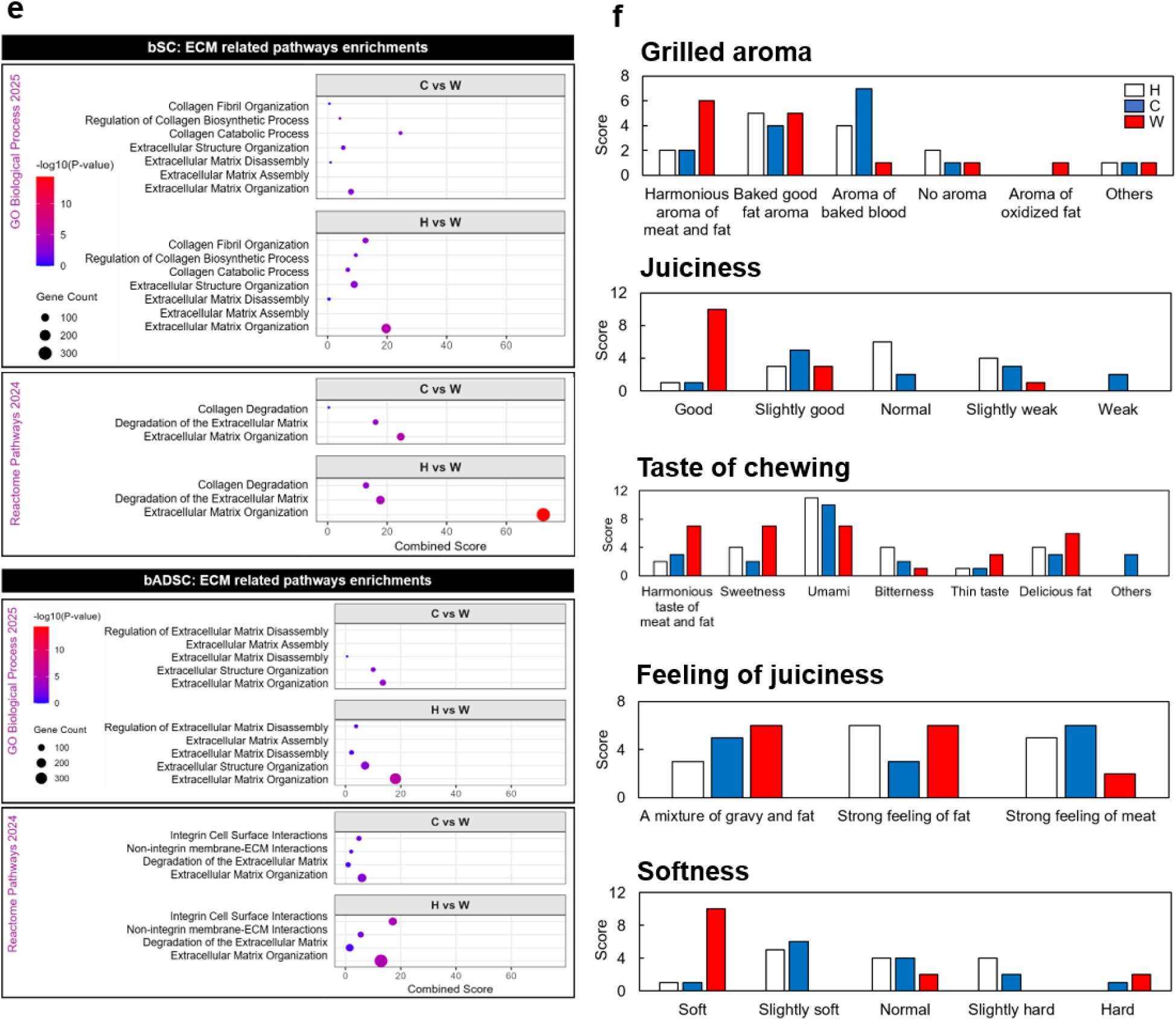
RNA sequence and sensory analyses of each meat. **a**, PCA analyses of RNA-seq data of bovine satellite cells (bSC) and adipose derived stem cells (bADSC) extracted from each beef meat. **b**, Volcano plots of C vs. W and H vs. W of bSC and bADSC, respectively. The threshold of significant up and down regulation is q < 0.05 and |log2(fold change)| > 0.58. Gene ontology (GO) analysis of bSC and bADSC for myogenesis (**c**), adipogenesis (**d**) and ECM (**e**) related pathway enrichments in C vs. W and H vs. W, respectively. **f**, Sensory evaluation for 5 categories of each beef meat by 14 volunteers. Open, blue and red boxes denote H, C and W, respectively.

Gene Ontology (GO) enrichment analyses were performed to examine pathways related to myogenesis, adipogenesis, and extracellular matrix (ECM) organization in bSCs and bADSCs (Fig. 4c–e). Among the myogenesis-related pathways, muscle contraction and myofibril-associated pathways were more enriched in Crossbreed-derived bSCs than in Wagyu- and Holstein-derived bSCs. Holstein-derived bSCs showed greater enrichment of pathways associated with myotube differentiation and myosin filaments than the other groups. For adipogenesis-related pathways, those associated with preadipocytes, subcutaneous adipose tissue, and adipose tissue were more enriched in Holstein-derived bADSCs, followed by Crossbreed-derived cells. In the analysis of ECM-related pathways, including ECM organization, extracellular structure organization, ECM degradation, and collagen degradation, Holstein- and Crossbreed-derived bSCs and bADSCs showed similar enrichment patterns, with higher enrichment than in the corresponding Wagyu-derived cells. These results suggest that the transcriptomic profiles of bSCs and bADSCs differ among cattle types, with distinct patterns in pathways associated with muscle development, adipogenesis, and ECM organization.

Sensory evaluation was performed on rib roast samples of Wagyu, Crossbreed, and Holstein beef using 14 volunteers (Fig. 4f). The evaluation covered five categories. Wagyu received the highest scores for fat aroma in the grilled aroma category, fat juiciness in the juiciness category, sweet taste associated with fat in the chewing category, and juiciness and softness. Crossbreed ranked second in most categories, although it received the highest scores for some attributes, including baked blood aroma in the grilled aroma category and strong meat sensation in the juiciness category. Holstein received scores comparable to or higher than the other beef types for baked fat aroma and strong fat sensation, and the highest scores for umami and bitterness in the chewing category. These results demonstrate differences in sensory properties among the three types of conventional beef. Sensory evaluation was not performed on the cultivated fibers.

### Fabrication of cultivated muscle and fat fibers and analysis of physical properties and amino acid profiles

To investigate whether breed-associated characteristics of conventional beef were reflected in cultivated fibers, muscle and fat fibers were fabricated by 3D bioprinting using bSCs and bADSCs isolated from Wagyu, Crossbreed, and Holstein cattle, respectively. These fibers were designed as the muscle and fat components of cultivated meat, as previously reported^4^. A customized 3D printer equipped with 48 nozzles was used to print bSCs or bADSCs together with collagen microfibers (CMFs), which supported cell adhesion and alignment. Fibrinogen and thrombin were also used to form a fibrin gel that connected the printed fibers into structures with a diameter of approximately 1 mm (Fig. 5a). After differentiation in the respective media, muscle and fat fibers were obtained (Fig. 5b). A representative image of the fabricated muscle fibers is shown in Fig. 5c.

**Fig. 5.**
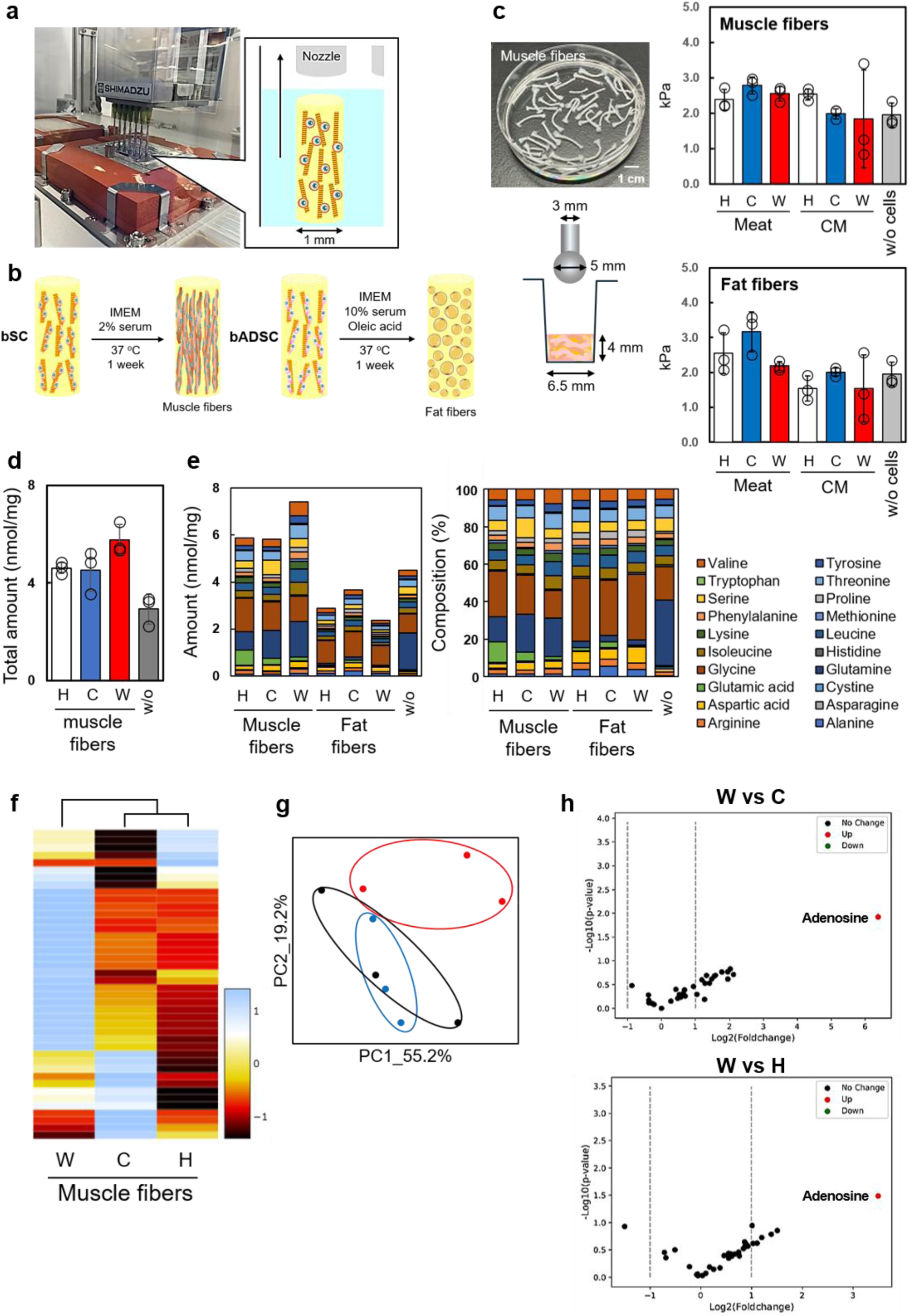
Fabrication of muscle and fat fibers differentiated from bSC and ADSC of each beef meat and amino acid analysis of these fibers. a, Picture and schematic illustration of customized 3D printer with 48 nozzles for cell fiber formation. b, Illustration of differentiation processes of bSC and ADSC fibers to muscle and fat fibers. c, Picture of the fabricated muscle fibers using Wagyu bSC. Illustration of compression experiments and the data of muscle and fat fibers of each beef meat, respectively (n=3). d, Molar amount of protein in muscle fibers constructed of bSC from each beef lean meat, respectively (n=3). e, Amount and composition ratio of free amino acids in muscle and fat fibers of each beef meat, respectively (n=3). f, Heat map analysis of free amino acids in muscle fibers from each beef lean meat. PCA analysis (g) and Volcano plots (h) of free amino acids in H vs. W and C vs. W., respectively.

The mechanical properties of the cultivated muscle and fat fibers were evaluated by compression testing (Fig. 5c). The elastic moduli of the cultivated fibers were approximately 70% of those of the corresponding native beef tissues. Nevertheless, the relative trends among the three cattle types were generally similar to those observed in native beef tissues, particularly for the fat portions.

The free amino acid profiles of the cultivated muscle fibers are shown in Fig. 5d. No statistically significant differences were observed among fibers derived from the three cattle types, although the overall profiles showed some similarities to those of the lean portions of conventional beef (Fig. 2d). Notably, Wagyu-derived muscle fibers showed the highest total free amino acid content (72.8 nmol/mg), following the same trend as that observed in the corresponding conventional beef. The amounts of amino acids detected from the cultivated muscle fibers were approximately one-third of those detected from the lean portions of conventional beef, whereas the amounts detected from the cultivated fat fibers were approximately comparable to those from the native fat portions in Fig. 2f (Fig. 5e). Glycine, which contributes to sweetness, was frequently detected in both cultivated muscle and fat fibers, as well as in the lean and fat portions of conventional beef. In contrast, alanine, another amino acid associated with sweetness, was detected less frequently in both types of cultivated fibers.

Heat map analysis of free amino acids in the cultivated muscle fibers showed that Crossbreed- and Holstein-derived fibers had similar profiles, whereas Wagyu-derived fibers showed a distinct profile (Fig. 5f). This pattern differed from that observed in conventional beef, in which Wagyu and Crossbreed were more similar to each other than to Holstein (Fig. 2e). PCA of the free amino acid profiles in the cultivated muscle fibers showed a similar pattern, with Crossbreed and Holstein samples distributed close to each other and separated from Wagyu samples (Fig. 5g). Volcano plot analysis of free amino acids, nucleosides, and vitamins in the cultivated muscle fibers showed that adenosine was less abundant in Wagyu-derived fibers than in Crossbreed- and Holstein-derived fibers, consistent with the trend observed in the lean portions of conventional beef (Fig. 5h). These results indicate that some breed-associated trends in amino acid profiles were reflected in the cultivated fibers, although the overall profiles were not identical to those of the corresponding native beef tissues.

### Fatty acid and aroma profiles of cultivated muscle and fat fibers

The fatty acid and aroma profiles of the cultivated fibers were compared with those of the corresponding native beef tissues. The total fatty acid content of the cultivated fat fibers derived from bADSCs was approximately one-hundredth of that in the original fat portions of conventional beef, although the relative trend among the three cattle types was similar, with Wagyu-derived fat fibers showing the highest value (Figs. 6a and 3a). The proportions of SFAs and MUFAs in the cultivated fat fibers were approximately 40–50%, similar to those in the native fat portions (Fig. 6b). In contrast, the PUFA proportion was approximately 30%, nearly ten times higher than that in the native fat portions.

**Fig. 6.**
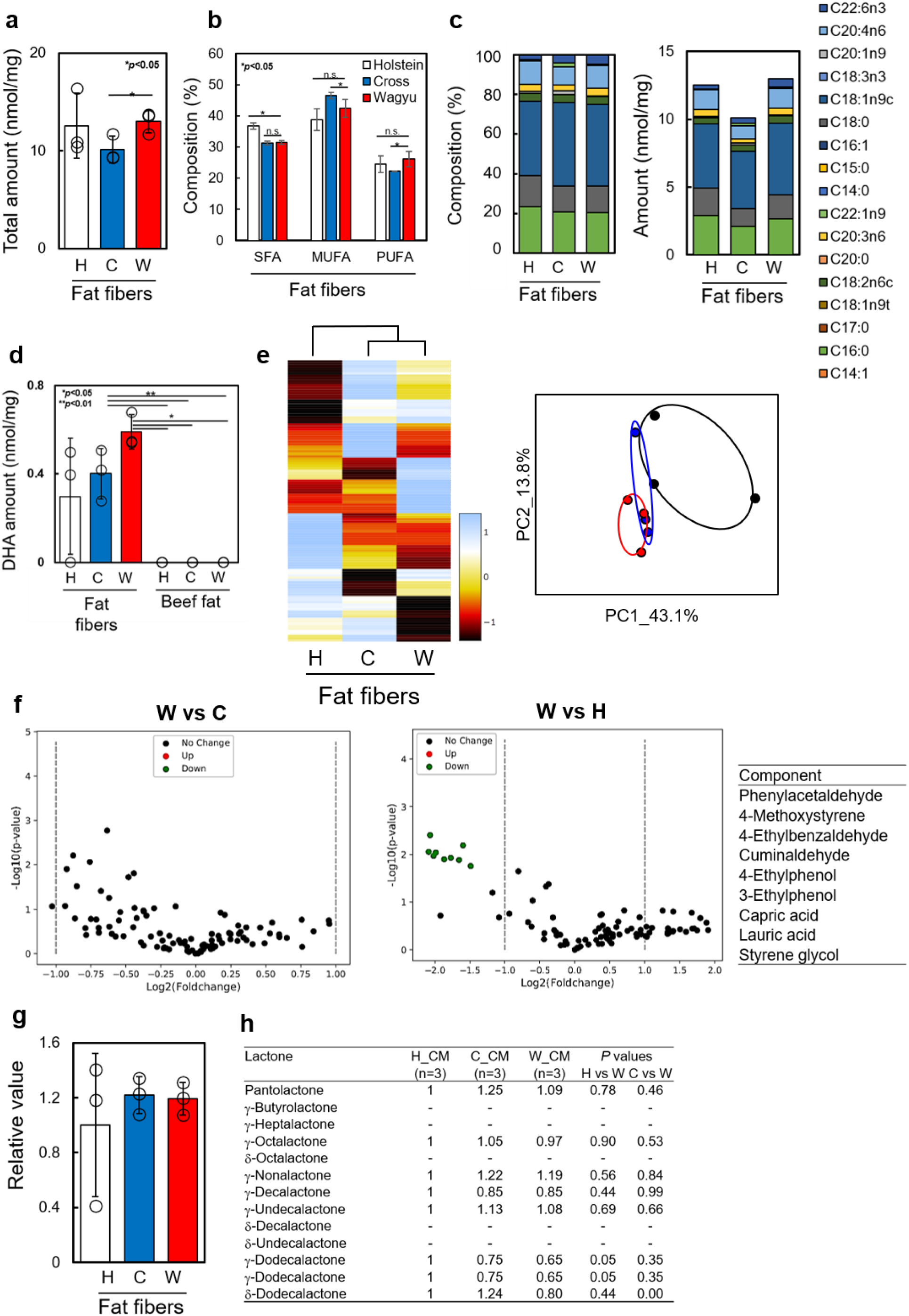
Fatty acid and aroma analyses of muscle and fat fibers. **a**, Total amount of fatty acids in fat fibers from each beef meat (n=3). **b**, Composition of SFAs, MUFAs and PUFAs in the fat fibers (n=3). **c**, Composition of fatty acids in the fat fibers. **d**, DHA amount in the fat fibers and the original fat tissues (n=3). **e**, Heat map and PCA analysis of aroma components in the fat fibers. **f**, Volcano plots of aroma components of grilled fat fibers in between W vs. H (left) and W vs. C (right), respectively. Summary of the aroma compounds with significant difference between W vs. H, fold change>1.5 (right table). **g**, Relative value of γ-nonalactone of fat fibers from each beef meat w/ heating. **h**, Summary of the relative values of the secreted lactone species from the grilled fat fibers in W and C as compared to those from H, respectively.

The profiles of individual fatty acids in the cultivated fat fibers showed some similarities to those of the native fat portions. Oleic acid was the predominant fatty acid in all cultivated fat fibers, followed by palmitic acid and stearic acid (Fig. 6c). The proportion of oleic acid was highest in Wagyu-derived fibers (43%), followed by Crossbreed (42%) and Holstein (37%). The proportions of palmitic acid and stearic acid were similar among the three groups, at approximately 20–23% and 15–18%, respectively. A notable difference between the cultivated fat fibers and native fat portions was the higher proportion of arachidonic acid (C20:4n6) in the cultivated fibers (10–15%), which has been reported to contribute to the taste of foods17. Furthermore, docosahexaenoic acid (DHA, C22:6n3) was detected in all cultivated fat fibers, with the highest amount observed in Wagyu-derived fibers (approximately 5% of total fatty acids, or 0.6 nmol/mg). In contrast, DHA was below the detection limit in the original fat portions (Fig. 6d).

Heat map and PCA analyses of aroma compounds released from the cultivated fat fibers showed that Wagyu- and Crossbreed-derived fibers had relatively similar profiles, whereas Holstein-derived fibers showed a somewhat different profile (Fig. 6e). This pattern was broadly consistent with the aroma profiles of the corresponding native fat portions (Fig. 3d–g). Volcano plot analysis of aroma compounds released from grilled cultivated fat fibers revealed that nine compounds, including lauric acid, were more abundant in Wagyu-derived fibers than in Holstein-derived fibers (Fig. 6f). The relative amount of γ-nonalactone released from Wagyu- and Crossbreed-derived fat fibers was higher than that from Holstein-derived fibers (Fig. 6g). The relative amounts of lactone species released from grilled cultivated fat fibers are summarized in Fig. 6h. Wagyu- and Crossbreed-derived fibers showed higher values for some lactones, including pantolactone, γ-nonalactone, and γ-undecalactone. However, this pattern differed from that of the native Wagyu fat portions, which showed the highest values for all lactone species (Fig. 3j). These results indicate that some breed-associated differences in fatty acid and aroma profiles were reflected in the cultivated fat fibers, although the amounts and relative compositions of these compounds differed from those in native beef tissues.

### Transcriptomic and proteomic profiles of cultivated muscle and fat fibers

The gene and protein expression profiles of the cultivated muscle and fat fibers were analyzed by RNA-seq and proteomic analysis, respectively (Fig. 7 and Supplementary Fig. S4). PCA of the RNA-seq data showed that Wagyu- and Crossbreed-derived fibers had relatively similar expression profiles in both muscle and fat fibers, whereas Holstein-derived fibers were more distinct (Fig. 7a). No apparent clustering attributable to batch effects was observed in the PCA plots. Volcano plots comparing Wagyu with Crossbreed or Holstein are shown in Fig. 7b. In cultivated muscle fibers, the numbers of significantly upregulated and downregulated genes in Crossbreed relative to Wagyu were 2,077 and 1,123, respectively, whereas the corresponding numbers in Holstein relative to Wagyu were 1,332 and 1,710. In cultivated fat fibers, the numbers of upregulated and downregulated genes were lower: 957 and 385 for Crossbreed relative to Wagyu, and 844 and 740 for Holstein relative to Wagyu.

**Fig. 7.**
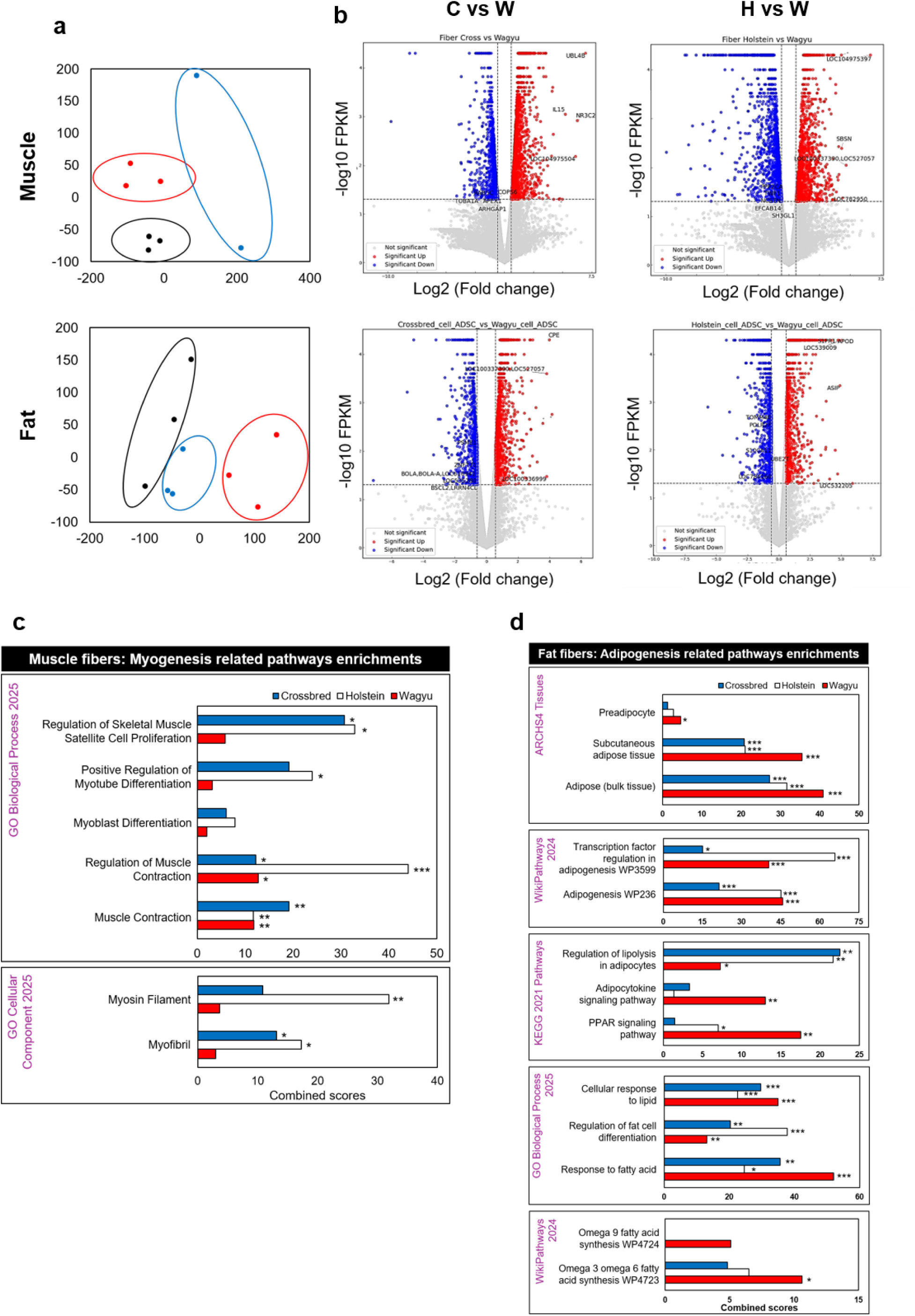
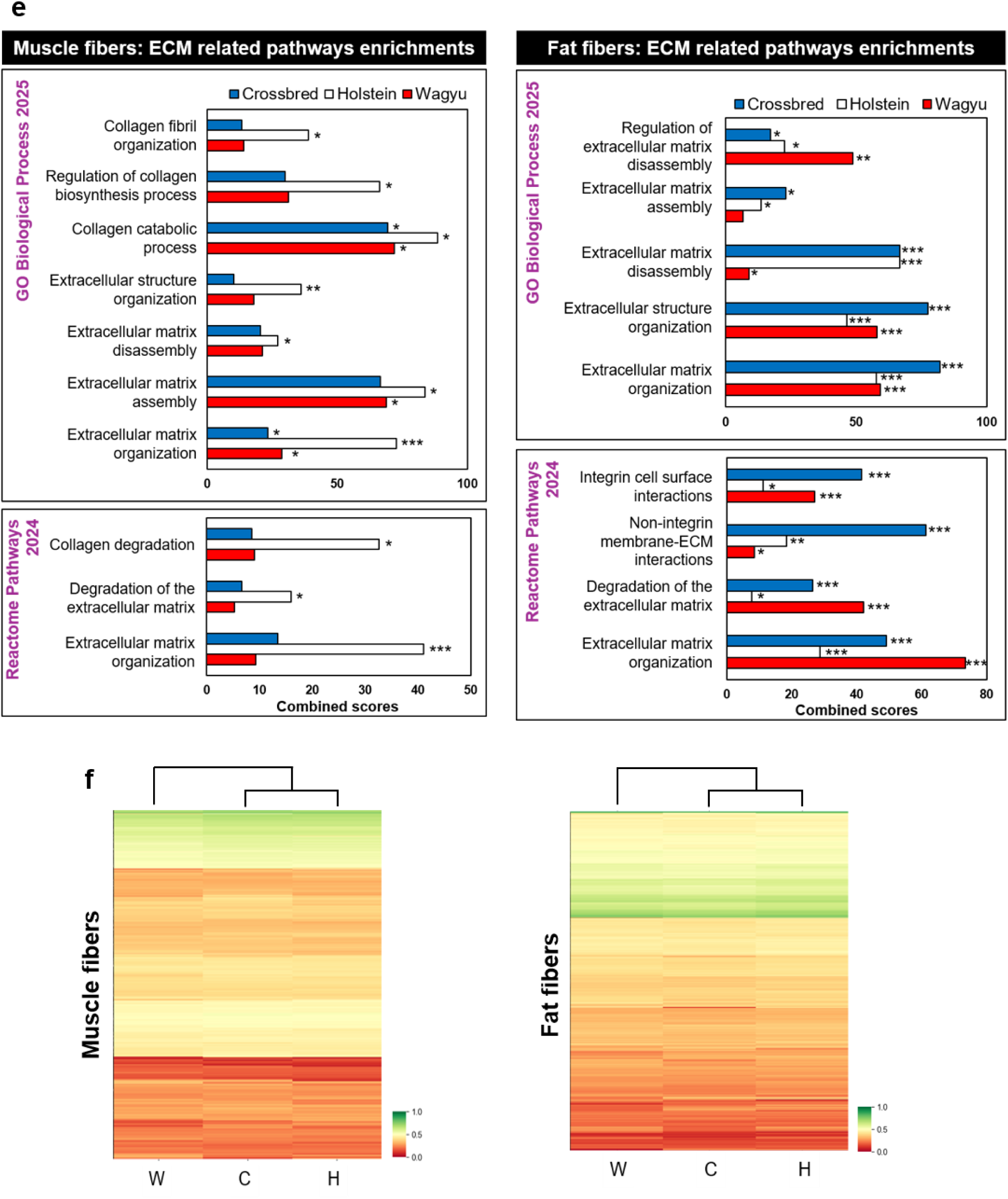
RNA sequence and protein expression analyses of muscle and fat fibers. **a**, PCA of RNA-seq data of muscle and fat fibers from each beef meat. **b**, Volcano plots of RNA-seq data of C vs. W and H vs. W of muscle and fat fibers, respectively. The threshold of significant up and down regulation is q < 0.05 and |log2(fold change)| > 0.58. GO analysis of muscle and fat fibers for myogenesis (**c**), adipogenesis (**d**) and ECM (**e**) related pathway enrichments in C vs. W and H vs. W, respectively. Asterisk (*) denotes significant statistical enrichments (\**p*<0.05, ** *p*<0.01 and *** *p*<0.001). **f**, Heat map protein expression analysis of muscle and fat fibers from each beef meat.

GO enrichment analyses of differentially expressed genes in the cultivated muscle and fat fibers, comparing Crossbreed with Wagyu and Holstein with Wagyu, revealed distinct enrichment patterns of myogenesis-associated processes in Holstein- and Crossbreed-derived fibers than in Wagyu-derived fibers (Fig. 7c). These results are consistent with the stronger myogenic gene expression profiles observed in Holstein- and Crossbreed-derived muscle fibers. Previous studies have reported differences in muscle fiber characteristics among cattle breeds, including a greater proportion of fast-twitch glycolytic fibers in Holstein cattle than in Wagyu cattle, which are characterized by relatively oxidative muscle fibers^18^. The enrichment of myogenesis-related pathways in Holstein- and Crossbreed-derived fibers may therefore reflect breed-associated differences in muscle development and contractile properties.

In contrast, Wagyu-derived fat fibers showed greater enrichment of adipogenesis-related pathways than Holstein- and Crossbreed-derived fibers (Fig. 7d). Pathways associated with adipocyte differentiation and lipid metabolism were particularly enriched in Wagyu-derived samples. This pattern is consistent with the high intramuscular fat deposition and fine marbling characteristic of Wagyu beef. Activation of the PPARγ regulatory network is a hallmark of adipocyte differentiation and lipid accumulation, and its upregulation has been associated with enhanced lipogenic activity and monounsaturated fatty acid synthesis in Wagyu cattle^19^. The enrichment of adipogenesis-related pathways in Wagyu-derived fat fibers suggests that the cells retain distinct adipogenic characteristics associated with their breed of origin. In addition, the enrichment of omega-6 and omega-9 fatty acid synthesis pathways in Wagyu-derived fibers suggests differences in fatty acid metabolism that may contribute to the breed-associated fatty acid profiles observed in the cultivated fibers.

ECM-related processes were also enriched in both cultivated muscle and fat fibers, with distinct patterns among the cattle types (Fig. 7e). In muscle fibers, Holstein-derived samples showed enrichment of pathways related to collagen fibril organization, ECM assembly, and collagen catabolism, suggesting differences in ECM remodeling during muscle fiber development. In contrast, Wagyu-derived fat fibers showed greater enrichment of pathways associated with ECM disassembly, integrin–cell surface interactions, and ECM–receptor interactions. ECM remodeling is involved in adipocyte expansion and tissue development during lipid accumulation^20^. These differences suggest that breed-associated characteristics are reflected not only in muscle- and adipogenesis-related pathways but also in ECM-related gene expression profiles.

Taken together, the transcriptomic analyses revealed distinct breed-associated expression profiles in cultivated muscle and fat fibers. Holstein-derived muscle fibers showed stronger enrichment of myogenesis- and ECM-related pathways, whereas Wagyu-derived fat fibers showed stronger enrichment of adipogenesis- and lipid metabolism-related pathways. These findings provide molecular evidence that some characteristics associated with the original cell sources are reflected in the cultivated fibers. Proteomic analysis further revealed differences in protein expression profiles among the cultivated muscle and fat fibers (Fig. 7f). Overall, the RNA and protein expression profiles of the cultivated fibers differed among the three cattle types, with Wagyu- and Crossbreed-derived fibers showing relatively similar profiles in several analyses.

## Discussion

The organoleptic properties of CM are considered to be affected by its composition (moisture, proteins, amino acids, fatty acids, lipids, vitamins, and food additives)^21^, production method, scaffold type and components, and post-processing^9,22^. Therefore, comparing the properties of CM samples reported in different studies remains challenging because their compositions, scaffolds, production methods, and cellular components vary considerably. In particular, when different cell types are used, even within the same species, it is difficult to determine whether differences in the resulting CM are attributable to cell origin or to the fabrication process. To address this issue, we systematically compared beef from Wagyu, Crossbreed, and Holstein cattle with cultivated muscle and fat fibers fabricated from cells derived from these cattle using the same 3D bioprinting method. This approach enabled us to investigate the extent to which breed-associated characteristics are reflected in cultivated fibers.

We first evaluated the physicochemical and biological characteristics of conventional beef from the three cattle groups. Texture profile analysis^23^ indicated that Wagyu beef was softer than Crossbreed and Holstein beef in terms of hardness, chewiness, and cohesiveness after boiling, although no corresponding difference was observed in springiness (Fig. 2a,b). Previous studies have reported weaker mechanical properties in Wagyu beef, potentially associated with remodeling of intramuscular connective tissue (IMCT)^24^. The H&E-stained sections of the beef samples were also consistent with differences in tissue architecture (Supplementary Fig. S1a,b). Because springiness reflects the recovery response during repeated compression and is related to the viscoelastic properties of a material^23^, the observed differences may also reflect differences in the relative contributions of connective tissue and adipose tissue among the beef samples. Owing to the limited size of the cultivated fibers, we used conventional compression testing rather than texture profile analysis to evaluate their mechanical properties (Fig. 5c). The elastic moduli of the cultivated muscle and fat fibers were approximately 70% of those of the corresponding native beef tissues. Nevertheless, the relative differences among the three cattle groups showed a similar trend, with Crossbreed fibers exhibiting the highest modulus, followed by Holstein and Wagyu fibers. These findings suggest that some breed-associated differences in mechanical properties were reflected in the cultivated fibers, although their mechanical properties did not fully reproduce those of native beef tissues.

Analysis of nutrient and free amino acid compositions showed that Wagyu beef contained the highest lipid content in both lean and fat portions, as well as relatively high levels of several free amino acids, including amino acids associated with sweetness, such as ornithine and alanine, and histidine, which has been associated with bitter or bittersweet taste (Fig. 2c–g). The higher lipid content in the lean portion of Wagyu beef is consistent with its characteristic intramuscular fat deposition. Intramuscular fat is known to contribute to beef quality, including juiciness, flavor, and tenderness^25^. Previous studies have also reported higher concentrations of monounsaturated fatty acids (MUFAs) in Wagyu beef than in beef from Holstein, Brown, and Charolais cattle^26^. In particular, Wagyu beef has been reported to contain higher proportions of fatty acids such as palmitoleic acid (C16:1), oleic acid (C18:1), and eicosenoic acid (C20:1). Oleic acid has also been associated with the characteristic aroma of Wagyu beef^27^. Our results showed similar trends in fatty acid composition, with Crossbreed beef generally exhibiting intermediate characteristics between Wagyu and Holstein. In addition, lactones, which have been reported to contribute to the characteristic sweet aroma of roasted Wagyu beef^28^, were present at higher levels in Wagyu fat tissue than in the other groups (Fig. 3j). Together, these results confirmed breed-associated differences in the physicochemical and compositional properties of the conventional beef samples examined in this study.

The cultivated muscle and fat fibers exhibited some compositional trends similar to those observed in the corresponding native beef tissues. However, their overall composition differed from that of native tissues. The free amino acid content of cultivated muscle fibers showed a trend similar to that of native lean meat, with the highest value observed in Wagyu-derived fibers, although the differences among the cultivated fibers were not statistically significant. The amount of free amino acids measured in the cultivated muscle fibers was approximately one-third of that in native lean meat, whereas the corresponding amount in cultivated fat fibers was comparable to that in native fat tissue (Fig. 5d,e). The major free amino acids detected in the cultivated fibers included glycine, glutamine, glutamic acid, serine, leucine, and valine, broadly reflecting the profiles of native beef tissues. However, alanine accounted for a lower proportion in the cultivated fibers than in native beef. Because alanine and glycine are associated with sweetness in beef, these compositional differences may influence the taste characteristics of cultivated fibers. Consistent with this possibility, Hwang et al. reported differences in taste characteristics between cultivated muscle tissues and native meat from chicken and cattle using an electronic tongue system, including lower umami, bitterness, and sourness values in the cultivated tissues^29^. The differences observed here may partly be related to the relatively short differentiation period of the cultivated fibers compared with the long growth period of cattle.

The fatty acid profiles of cultivated fat fibers also showed both similarities to and differences from those of native fat tissues. The proportions of SFAs and MUFAs were broadly similar between cultivated fat fibers and native fat tissues, whereas the proportion of PUFAs was approximately ten times higher in the cultivated fat fibers (Fig. 6c). Oleic acid was the most abundant fatty acid in all cultivated fat fibers, followed by palmitic and stearic acids, as observed in native fat tissues. Notably, DHA was detected in the cultivated fat fibers, whereas it was not detected in the native fat tissues under the conditions used in this study (Fig. 6d). Although the expression of fatty acid desaturase genes was higher in the corresponding ADSC fibers than in the differentiated fat fibers (Supplementary Table S1), the relationship between these expression patterns and DHA accumulation requires further investigation. In addition, the total fatty acid amount in cultivated fat fibers was approximately one-hundredth of that in native fat tissue. These results indicate that, although some aspects of the fatty acid composition were similar, the amount and relative composition of fatty acids in cultivated fat fibers were not yet equivalent to those in native tissues.

Gene expression analyses further revealed differences associated with the cellular origins of the cultivated fibers. In particular, Wagyu-derived fat fibers showed higher expression of genes and enrichment of pathways related to adipogenesis and lipid metabolism than those derived from Crossbreed and Holstein cattle. These findings are consistent with the characteristic intramuscular fat deposition of Wagyu beef and suggest that some adipogenic features associated with the cell source are retained during in vitro differentiation. Previous studies have reported associations between polymorphisms in the mitochondrial transcription factor A (TFAM) gene and fat deposition traits, including marbling, in cattle^30^. Mitochondrial function is also closely associated with adipocyte metabolism, and brown adipose tissue (BAT) is characterized by high mitochondrial activity and energy expenditure^31^. Although there is no definitive evidence that Wagyu cattle have a greater amount or higher activity of BAT than other cattle breeds, we examined the expression of genes associated with BAT, including uncoupling protein 1 (UCP1), which mediates mitochondrial uncoupling and heat production^32^. The expression of UCP1 in Wagyu-derived fat fibers was 1.625-fold higher than that in Crossbreed-derived fibers (Supplementary Table S2). We also examined the expression of PR domain containing 16 (PRDM16), a regulator associated with brown adipocyte differentiation^33^. The expression of PRDM16 was 1.88-fold higher in Wagyu-derived fat fibers than in Crossbreed-derived fibers (Supplementary Table S2). These results demonstrate that Wagyu-derived fat fibers exhibit higher expression of BAT-associated genes than Crossbreed-derived fibers, although further studies are required to determine whether these differences are accompanied by functional characteristics of brown adipocytes. In contrast, Holstein-derived muscle fibers showed enrichment of pathways associated with myogenesis and extracellular matrix (ECM) organization. These differences suggest that the cellular origin may influence the molecular characteristics of cultivated muscle and fat fibers, even when the same fabrication and differentiation procedures are used.

This work provides a systematic comparison of the physicochemical and biological properties of beef from Wagyu, Wagyu–Holstein Crossbreed, and Holstein cattle, together with those of cultivated muscle and fat fibers fabricated from cells isolated from these cattle. The overall trends in breed-associated characteristics observed in conventional beef and the corresponding cultivated fibers are schematically summarized in Fig. 8. The comparisons of physical properties, amino acid composition, protein profiles, fatty acid composition, and flavor-related characteristics revealed similarities in some breed-associated trends between conventional beef and the corresponding cultivated fibers. Although the cultivated fibers did not fully reproduce the properties of native beef tissues, these findings suggest that some characteristics associated with the source breed can be reflected in cultivated muscle and fat fibers. In addition to these physicochemical and compositional findings, the GO and Hallmark gene set-based clustering analyses revealed distinct gene expression patterns among the three breeds in both the undifferentiated cells (bSCs and bADSCs) and the corresponding differentiated muscle and fat fibers (Supplementary Figs. S5 and S6). These findings suggest that breed-associated molecular characteristics are present in the source cells and remain, at least in part, reflected in the gene expression profiles of the differentiated fibers. Although the expression patterns were not identical before and after differentiation, the results support the contribution of cellular origin to the molecular characteristics of cultivated tissues. For example, Wagyu-derived cells exhibited higher expression of genes associated with adipogenesis and lipid metabolism, whereas Holstein-derived cells showed enrichment of pathways related to myogenesis and extracellular matrix organization. Dendrograms of bSC, muscle fibers, bADSC, and fat fibers represented that original cell characters might be retained in the differentiated fibers (Supplementary Fig. S7). Together, these results suggest that selecting cells from different cattle breeds may provide a means of tailoring the biological and compositional properties of cultivated muscle and fat tissues. Nevertheless, differences in the properties of cultivated fibers and native beef tissues remain. For example, the total fatty acid content of the cultivated fat fibers was substantially lower than that of native fat tissue, despite similarities in the relative proportions of some fatty acid classes. Differences were also observed in the fatty acid profiles, including the detection of DHA in cultivated fat fibers. These findings indicate that the properties of cultivated fibers are influenced not only by cellular origin but also by the in vitro differentiation environment. In this regard, our previous study demonstrated that the fatty acid composition of cultivated fat derived from Wagyu bADSCs could be modulated by changing the matrix components during differentiation, resulting in fat in which oleic acid accounted for more than 80% of the MUFAs^34^. This finding highlights the potential to tailor the fatty acid composition of cultivated fat through the optimization of the culture environment. Together, these results suggest that combining the selection of cell sources with the optimization of differentiation conditions and matrix components may enable more precise control over cultivated meat properties. Thus, this study provides a basis for developing cultivated meat with selected breed-associated characteristics, although further optimization of cell differentiation and tissue fabrication will be required to more closely reproduce the physicochemical and compositional properties of conventional beef.

**Fig. 8.**
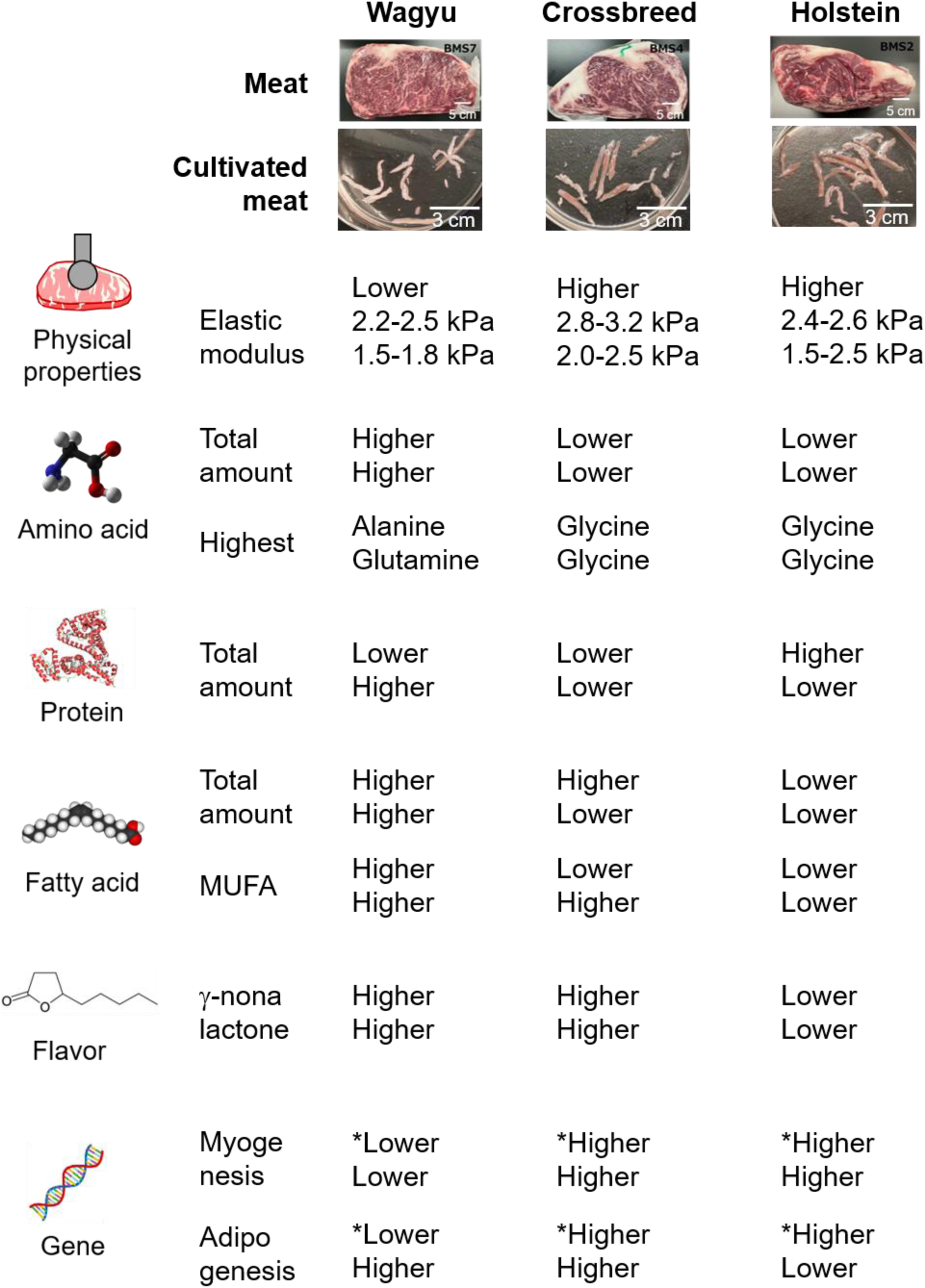
Summary of the analyses and characteristic comparison between each beef meat and cultivated meat. Biological, physicochemical and sensory analyses of W, C and H, respectively. Asterisk denotes that gene expressions of bSC and bADSC before differentiation to myotube and adipocyte, respectively.

## Methods

Unless otherwise stated, all reagents were of analytical grade and used without further purification. 2-Morpholinoethanesulfonic acid monohydrate (used as an internal standard) was obtained from Dojindo Laboratories. Methanol, chloroform, amino acids mixture standard solutions Type AN and Type B, L-asparagine standard solution, L(+)-glutamine reference material, L-tryptophan reference material), adenosine, and 0.1% formic acid in acetonitrile (Cat# 068-04723) were all purchased from FUJIFILM Wako Pure Chemical Corporation. Adenosine 5′-monophosphate disodium salt (Cat# A2117) was obtained from Tokyo Chemical Industry Co., Ltd. (TCI). LC-MS grade 0.1% formic acid in water (Cat# LS118-4) was purchased from Fisher Chemical, and 2-propanol (Cat# 29112-63) was obtained from Nacalai Tesque, Inc. Hexane for pesticide residue and polychlorinated biphenyl (PCB) tests (Cat# 083-07911) was purchased from FUJIFILM Wako Pure Chemical Corporation. The Supelco 37 Component FAME Mix was purchased from Sigma-Aldrich. Palmitic acid-d₃₁ was obtained from Cambridge Isotope Laboratories, Inc. A fatty acid methylation kit was obtained from Nacalai Tesque. Methanol, chloroform, acetone, and hexane for pesticide residue and PCB tests were all purchased from Kanto Chemical Co., Inc. Distilled water used for GC–MS analysis was pre-washed with hexane (Kanto Chemical). Dulbecco’s phosphate-buffered saline without calcium and magnesium was purchased from Nacalai Tesque, Inc. A 4% paraformaldehyde phosphate buffer solution was purchased from FUJIFILM Wako Pure Chemical Corporation.

### Texture Profile Analysis

Texture profile analysis was performed using an EZ-TEST^TM^ (SHIMADZU) equipped with a 500 N load cell (Shimadzu, Kyoto, Japan). Rectangular samples measuring 3 cm × 2.5 cm × 1.0 cm were prepared by cutting along the muscle fibers. Prior to analysis, the samples were immersed in a water bath maintained at 70°C for 1 hour, followed by cooling under running water for 30 minutes, and then equilibrated to room temperature (approximately 25°C) for at least 30 minutes. Each sample was subsequently subjected to a double compression test, during which the sample was compressed to 50% of its original height at a test speed of 1.0 mm/s. From the resulting force–time curves, textural parameters including hardness, cohesiveness, springiness, gumminess, and chewiness were calculated. All measurements were conducted (n = 3), and the mean values were used for subsequent statistical analysis.

### Compression Elasticity Measurement

The compression elasticity of the samples was measured using an EZ-TEST (Shimadzu, Kyoto, Japan) equipped with a 20 N load cell. All fibers—including fibers of cultivated meat and fibrous tissues derived from conventional meat—were washed by PBS several times prior to measurement. After preparation, the fibers were stacked in a 24-well insert so as to completely cover the bottom surface of the insert. A spherical mold (5 mm in diameter) was employed to measure the elastic modulus at a head-moving speed of 1.0 mm/min. The compressive test protocol involved gradually increasing the engineering strain until the applied stress reached 200 mN. The elastic modulus was automatically calculated by the software TRAPEZIUM^TM^ X-V (SHIMADZU) within the elastic range (10–20 mN). The total sample size was n = 3 for each fiber type. Due to the difficulty in directly measuring the sample height, the height was defined as the elongation measured at a compression force of 200 mN, corresponding to the point at which sufficient force was applied to the lower compression plate.

### Proximate Composition Analysis

The analysis of moisture, protein, lipid, ash, carbohydrate, and energy content was outsourced to the Japan Food Research Laboratories (General Incorporated Foundation). The following methods were applied. Approximately 100 g of each solid sample was submitted for analysis.

#### Moisture

Determined by atmospheric heating-drying method or vacuum heating-drying method.

#### Protein

Measured by the combustion method.

#### Lipid

Analyzed using the Soxhlet extraction method or acid decomposition method.

#### Ash

Determined by the direct ashing method.

#### Carbohydrate

Calculated by difference as: 100 – (moisture + protein + lipid + ash).

#### Energy

Calculated using the following energy conversion factors: protein, 4 kcal/g; lipid, 9 kcal/g; carbohydrate, 4 kcal/g.

### Amino acid analysis

Frozen meat samples (separated into lean and fat) and cultivated meat samples were ground under dry ice conditions and subsequently stored at -80°C. One cultivated meat fiber with measured weight (later converted) and about 10 mg of the ground meat sample were used. Following the addition of 10 µL of an internal standard solution (0.5 mg/mL 2-morpholinoethanesulfonic acid monohydrate), protein precipitation was performed using a mixed solvent composed of methanol, water, and chloroform in a 2.5:1:1 ratio. The supernatant was then collected, freeze-dried, and reconstituted in ultrapure water. Quantitative analysis of 20 free amino acids was conducted via LC-MS using calibration curves, while the remaining 141 species were relatively quantified based on the peak area ratios relative to the internal standard. LC-MS analyses were performed using LCMS-TQ^TM^ 8060NX (SHIMADZU). Separation was achieved on a column Discovery HS F5-3 (150 mm × 2.1 mm, 3 µm). The mobile phase consisted of solvent A (0.1% formic acid in water) and solvent B (0.1% formic acid in acetonitrile), delivered at a flow rate of 0.25 mL/min with the column maintained at 40°C. Gradient elution conditions were set in accordance with Smart Metabolites Database^TM^ (SHIMADZU). Detection was carried out in positive ion mode using electrospray ionization (ESI), monitoring the [M+H]⁺ ions for each amino acid. For multi-omics analyses, an integrated multi-omics analysis package including Smart Metabolites Database^TM^ (SHIMADZU) was employed to consolidate the results. After quantification of amino acids and nucleic acid-related metabolites, statistical analysis and data visualization were performed using LabSolutions Insight^TM^ and the Multi-omics Analysis Package (SHIMADZU). Volcano plots and heatmaps were generated using the software. In the volcano plot, metabolites with adjusted p-values less than 0.05 and absolute log2 fold changes greater than 1 were highlighted as significantly different. The heatmap visualized relative concentration differences of each metabolite with hierarchical clustering.

### Fatty acid analysis

Frozen meat samples separated into fat and cultivated meat samples were ground under dry ice conditions and subsequently stored at –80°C. One cultivated meat fiber with measured weight (later converted) and about 10 mg of the ground meat sample were used. Fatty acid composition was analyzed using gas chromatography-mass spectrometry (GC-MS). Palmitic acid-d₃₁ was methylated to its fatty acid methyl ester (FAME) and diluted with n-hexane to prepare an internal standard solution at a final concentration of 10 μg/mL. A calibration curve was constructed using standard solutions, and its linearity was confirmed (p > 0.9990). Sample solutions were appropriately diluted with chloroform, followed by methanolysis. The reaction mixture was subjected to liquid-liquid extraction, and the organic phase was collected for GC-MS analysis. GC-MS analysis was performed using a GCMS-QP^TM^ 2020 system (SHIMADZU) equipped with a TC-70 capillary column (60 m × 0.25 mm i.d., 0.25 μm film thickness; GL Sciences, J2102K05). The injector temperature was set at 280°C, and the injection volume was 1 μL in splitless mode. The oven temperature program was as follows: an initial temperature of 140°C was held for 1 min, followed by an increase at a rate of 3°C/min to 240°C, then further increased at 25°C/min to 280°C, where it was held for 5 min. Helium was used as the carrier gas at a flow rate of 25 cm/sec. The mass spectrometer was operated in electron ionization (EI) mode. The data acquisition range was not specified. Fatty acids were identified by comparing their retention times and mass spectra with those of authentic standards and spectral libraries such as NIST and Wiley. Quantification was performed using a calibration curve, and corrections were applied using the internal standard method. Fatty Acid Analysis was performed by Shimadzu Techno-Research, Inc. using their protocols.

### Aroma component analysis

Conventional and cultured meat samples were directly transferred into 20 mL headspace vials. An appropriate amount (about 20 mg) of each sample was transferred into a 20 mL headspace vial and sealed with a PTFE/silicone septum. For heated samples, the vials were placed on a heating plate set to 200°C for 30 minutes. After heating, the vials were allowed to cool to room temperature for a minimum of one hour prior to measurement. For raw (non-heated) samples, the sealed vials were kept at room temperature for at least one hour before measurement. Solid phase microextraction (SPME) and GC-MS analysis of aroma compounds was performed using multifunctional autosampler AOC^TM^-6000 and gas chromatograph-mass spectrometer GCMS-TQ^TM^ 8050 NX (SHIMADZU). An SPME Arrow (DVB/CarbonWR/PDMS, 20 mm x o.d. 1.1 mm) was employed for extraction. The vial was equilibrated at 80°C with agitation for 10 min, and then the SPME Arrow was exposed to the headspace of the vial for 30 min to adsorb the aroma compounds. The adsorbed volatiles were desorbed in the GC injector at 250°C in split mode (split ratio 1:2) and analyzed using GC-MS. GC separation was carried out on an InertCap PureWAX column (30 m × 0.25 mm inner diameter, 0.25 µm film thickness) (GL Sciences) with helium carrier gas at a constant pressure of 83.5 kPa. The following oven temperature was used: 50°C for 5 min, increased at 10°C/min to 250°C, and maintained for 10 min. Electron ionization (EI) was employed in the ionization mode, and data were acquired in scan mode (m/z 35 - 500). The compound annotation was performed based on the mass spectral and retention indices using Smart Aroma Database. Quantification of selected aroma compounds was performed using peak area normalization or external standard calibration to determine their relative concentrations. All analyses were conducted in triplicate to ensure reproducibility, and blank runs were performed to confirm the absence of background contamination. Statistical analysis and data visualization were performed using the LabSolutions Insight^TM^ and the Multi-omics Analysis Package (SHIMADZU). Volcano plots and heatmaps were generated using the software. In the volcano plot, metabolites with adjusted p-values less than 0.05 and absolute log2 fold changes greater than 1 were highlighted as significantly different. The heatmap visualized relative concentration differences of each metabolite with hierarchical clustering.

### RNA sequencing

RNA-seq analyses were performed using three biological replicates per breed (Wagyu, Crossbreed, and Holstein), each derived from an independent animal (n = 3 per breed). Specifically, cells derived from these animals were grown to confluence, and total RNA was extracted from the resulting cell pellets using the QIAwave RNA Mini Kit. For the 2D cultures, each sample consisted of 2 × 10^6^ cells. For the fiber cultures, RNA was extracted from a single 20 mm fiber that had undergone proliferation and differentiation and was then mechanically fragmented. Transcriptome analysis was outsourced to RIKEN Genesis Inc. (Kanagawa, Japan). For transcriptome sequencing, libraries were constructed using the TruSeq Stranded mRNA kit following the manufacturer’s protocol. Between 1 × 10^4^ and 1 × 10^5^ living cells per sample were used for library preparation. The sequencing was performed on the Illumina NovaSeq 6000 platform with paired-end sequencing, generating reads of 101 base pairs in length. On average, approximately 300 million reads per sample were obtained, with a total sequencing depth reaching 1.2 billion read pairs per run. The raw data were mapped to the iGenomes BOS taurus NCBI UMD_3.1.1 reference genome using Tophat for mapping, junction prediction, and fusion gene detection, followed by transcript assembly, expression quantification, and differential expression analysis using Cufflinks. FASTQ files were initially processed with PRINSEQ for quality trimming. For downstream analyses, genes with FPKM values in the lowest 25% were excluded.

Visualization steps included:

- Heatmap generation by the Python matplotlib library.
- Principal component analysis (PCA) by the sklearn package in Python.
- Volcano plots were constructed based on computed p-values (excluding entries with zero values) and fold-change values using matplotlib.

Comparative analyses were performed between Crossbred vs. Wagyu and Holstein vs. Wagyu for both cells and fibers. Differentially expressed genes (DEGs) were identified using the following thresholds:

q < 0.05

|log2(fold-change)| > 0.58

Gene ontology enrichment analysis was performed using the org.Bt.eg.db annotation library in R, with further analysis carried out using the clusterProfiler package and results visualized via ggplot. Further analyses of enriched pathways were performed using the Enrichr^35^ comprehensive gene set enrichment analysis web server (Ma’ayan Laboratory, Icahn School of Medicine at Mount Sinai; version updated October 17, 2025). Differentially expressed genes showing either upregulation or downregulation were selected and input as gene lists using their Entrez Gene symbols (approximately 1000 genes per comparison). Gene expression profiles of differentiated muscle fibers and fat fibers were compared between Wagyu and Crossbreed, and between Wagyu and Holstein, respectively. Functional enrichment was carried out using the GO Biological Process, KEGG, and WikiPathways libraries. Significantly enriched terms (adjusted p < 0.05) related to myogenesis, adipogenesis, and extracellular matrix (ECM) organization were then selected for comparative interpretation of gene expression among the three breeds.

### Sensory analysis

Ribeye (longissimus dorsi) samples were obtained from steers of three breeds: Japanese Black, Holstein, and Holstein–Japanese Black crossbred. (IH MEAT PACKER INC., Japan) All cattle were 20.9–29.5 months old. Carcasses were aged for 5 days postmortem, then frozen and stored at –20 °C. On the testing day, the muscle was thawed and sliced perpendicular to the fibers into uniform portions (5 cm × 0.8 cm), ensuring the thickness did not exceed 1 cm. Each piece was cooked on a preheated hot plate lined with aluminum foil and lightly greased with vegetable oil. Cooking was performed at 220 °C: 2 min on the first side, 2.5 min on the reverse, and 30 s on the sides to achieve even browning. The samples were rested under foil for 1 min before serving. Sensory evaluation was conducted by a panel of 14 inexperienced members. The attributes evaluated included grilled aroma, juiciness, feeling of juiciness, softness, and taste during chewing. Evaluations were performed under blind conditions, with sample presentation order randomized. Water was provided to cleanse the palate between samples. The study was approved by the Research Ethics Committee for Human Subject Studies, Graduate School of Engineering, Osaka University (Approval No. 4-1-1). Informed consent was obtained from all participants. Participants were recruited on a voluntary basis from within the research laboratory. No compensation was provided to the participants.

### Isolation and purification of bSC and bADSCs

Muscle and adipose tissues were obtained from the knuckle (shintama), located on the inner side of the round of Japanese Black cattle and crossbred cattle (a hybrid of Japanese Black and Holstein) at the slaughterhouse (Ariake Beef Plant of SANKYOMEAT INC., Japan). Tissues from Holstein cattle were obtained from a separate slaughterhouse (SANKYO Meat LTD. Ariake Meat Plant, Japan). The tissues were kept at 0–4 °C and carried to the laboratory after health inspection of the carcass. On the next day, bSCs and bADSCs were isolated from the respective tissues of each individual animal, basically according to the methods described previously^4,30^. To be handled in accordance with the “Food Sanitation Act” ⅲ), the tissue samples were disinfected using Perasan MP2-J (Enviro Tech Japan, Tokyo, Japan) at 0.18 % (w/v) as peracetic acid and the digestion was performed using trypsin (FUJIFILM Wako Pure Chemicals, Osaka, Japan) in place of collagenase. Cell sorting was replaced by the preplating method to increase the purity of bSCs. The 1st passage of bSCs and the 2nd passage of bADSCs were stored in 1 mL aliquots using Bambanker DMSO Free (NIPPON Genetics, Tokyo, Japan) (3 × 10^6^ cells/mL), respectively, according to the manufacturer’s instruction. Each lot was obtained from a different bovine donor. Accordingly, three independent lots derived from three individual animals were used for each breed (Wagyu, Crossbreed, and Holstein). No live animals were used or subjected to experimental procedures in this study; therefore, animal ethics approval was not required.

### 2D cell culture

bSCs were cultured in bSC proliferation medium containing high-glucose DMEM (Thermo Fisher Scientific, 10569-010) containing 1% antibiotic–antimycotic mixed solution (Nacalai tesque, 02892-54), 20% FBS (Thermo Fisher Scientific, 10270-106), 4 ng/mL basic fibroblast growth factor (bFGF) (FUJIFILM Wako Pure Chemical Corp Fujifilm, 067-04031), and 10μM p38 inhibitor (Selleck Chemicals, SB203580) for proliferation. bADSCs were cultured in ADSC proliferation medium containing high-glucose DMEM (Gibco, 10569-010) containing 1% antibiotic–antimycotic mixed solution (Nacalai tesque, 02892-54), 10% FBS.

### Fabrication of muscle and fat fibers by 3D printing

3D printing was performed using the tendon-inspired printing system (TIPS) method. TIPS is a technique in which the printing bath is vertically divided into three layers: a bottom tendon gel, a supporting bath, and an upper tendon gel, allowing for spatially controlled placement of printed structures. In this study, two bioinks were prepared separately: a bSC bioink containing bovine satellite cells at 5 × 10^7^ cells/mL, and an ADSC bioink containing adipose-derived stem cells at 5 × 10^6^ cells/mL. Each bioink also included collagen microfibers (CMF) at 1.2 wt % (12 mg/mL) and fibrinogen (Sigma-Aldrich, F8630) at a final concentration of 15 mg/mL. The supporting bath was prepared following Kang et al., using IMEM (Integriculture) containing 10% FBS and edible gelatin. The gelatin was sterilized by UV radiation for 45 minutes before being dissolved in the medium. Then gelatin bath was added to get a final concentration of 5 U/mL thrombin solution (Sigma-Aldrich, T4648) (the stock solution at 1,000 U/mL prepared in DMEM). Both the upper and lower tendon-gel layers were composed of a 4 wt% CNF solution prepared from type I collagen (Nippi, PSC-1-200-1000). Each fiber was produced by dispensing 70 µL of bioink with a custom-developed 48-nozzle multi-dispenser bioprinter (Shimadzu), enabling the simultaneous deposition of multiple constructs. After printing, the print cell containing the gelatin support bath was incubated at 25 °C for 30 minutes followed by 37 °C for 2 h to cure the fibrin gel. The liquefied gelatin was then removed and replaced with culture medium for proliferation using a peristaltic pump system. bSCs fibers were subjected to a 3-day proliferation phase followed by 4-day differentiation phase, while ADSCs fibers underwent a 3-day proliferation phase followed by a 14-day differentiation phase. bSCs were induced to differentiate in DMEM supplemented with 2% fetal bovine serum (FBS). ADSCs were differentiated in DMEM containing 10% FBS and 500 µM oleic acid (NOF Corporation). During differentiation, the bath medium (60 mL, corresponding to one-third of the total volume) was replaced every 8 h. All media used for fiber culture were further supplemented with 5 mM ε-aminocaproic acid (Sigma-Aldrich, A2504) and 1% (v/v) antibiotic– antimycotic solution.

### Water content analysis

The water content is calculated according to the mass before and after freeze-drying. Briefly, the printed fibers in PBS1x were taken out, the surface liquid was removed, and the wet weight (Wwet) of the fibers was measured by a balance. The dry weight (Wdry) of the fibers was measured after freeze-drying (24 h). The water content is given by the following formula (1): V_water_ = 100*W_wet_ – W_dry_ /W_wet_

### Histological analysis

Tissue samples were fixed in 4% neutral-buffered formalin at 4°C overnight. Paraffin embedding and sectioning at a thickness of 5 μm, as well as hematoxylin and eosin (HE) staining, were performed by Applied Medical Research using their standard protocols.

### Statistical analysis

Statistical analysis was performed using EzAnova (version 0.98, University of South Carolina, Columbia, SC, USA) software. The details of the number of n corresponding to the number of independent experiments using isolated bovine primary cells from different bovine donors, or independent samples, are displayed in the captions. A two-way ANOVA was applied with time set as “paired or repeated measures” and the treatment as classic analysis or “unpaired” which led to a pairwise comparison, with the Tukey’s HSD post hoc test for the multiple comparisons. Error bars represent SD. p values < 0.05 were considered to be statistically significant.

## Data availability

The data that support the findings of this study are available from the corresponding author upon reasonable request.

## Acknowledgements

The authors thank Shimadzu Analytical Innovation Research Laboratories. This study was financially supported by J231013003 from Ministry of Agriculture, Forestry and Fisheries (MAFF), JP22H05141, JP25H01220 and JP22K21348 from Japan Society for Promotion of Science (JSPS). This study was also supported by bilateral program (JPJSBP120252301) from JSPS, COI-NEXT (JPMJPF2009) from Japan Science and Technology Agency (JST), and 21W2K022, 23201377-0 and 23201347-0 commissioned by the New Energy and Industrial Technology Development Organization (NEDO).

## Author contributions

C.N., Y.K., K.S., A.T., K.N. and M.M. conceived of and designed the experiments.

C.N., F.L., A.Y., Miki M., T.S., K.N., and A.T. performed the experiments.

C.N., F.L., A.Y., T.S., K.N., and A.T. analyzed the data. C.N. and M.M. wrote the manuscript. All authors read and approved of the final manuscript.

## Competing interests

The authors declare that they have no competing financial interests.

**Supplementary information** is available for this paper at https://.

**Corresponding and requests for materials** should be addressed to M.M.

**Reprints and permission information** is available at www.nature.com/reprints.

## Publisher’s note

Springer Nature remains neutral with regard to jurisdictional claims in published maps and institutional affiliations.

