## Supplementaly information for "Breed-Specific Characteristics of Beef Are Partly Retained in Cultivated Meat"

**Contents of supporting information:**

**Fig. S1. Histological analysis and weight loss rate by heating**
**Fig. S2. Summary of secreted aroma components from grilled meats**
**Fig. S3. Phase contract image of bSC and bADSC**
**Fig. S4. RNA sequence analyses of muscle and fat fibers**
**Table S1. RNA expression of fatty acid desaturase 1 and 2**
**Table S2. RNA expression of UCP1 and PRDM16**
**Fig. S5. Clustering of gene expression profiles using GO and Hallmark gene sets in bSC and bADSC**
**Fig. S6. Clustering of gene expression profiles using GO and Hallmark gene sets in muscle and fat fibers**
**Fig. S7. Dendrograms of bSC, muscle fibers, bADSC, and fat fibers**
**Methods. Clustering of gene expression profiles using GO and Hallmark gene sets**

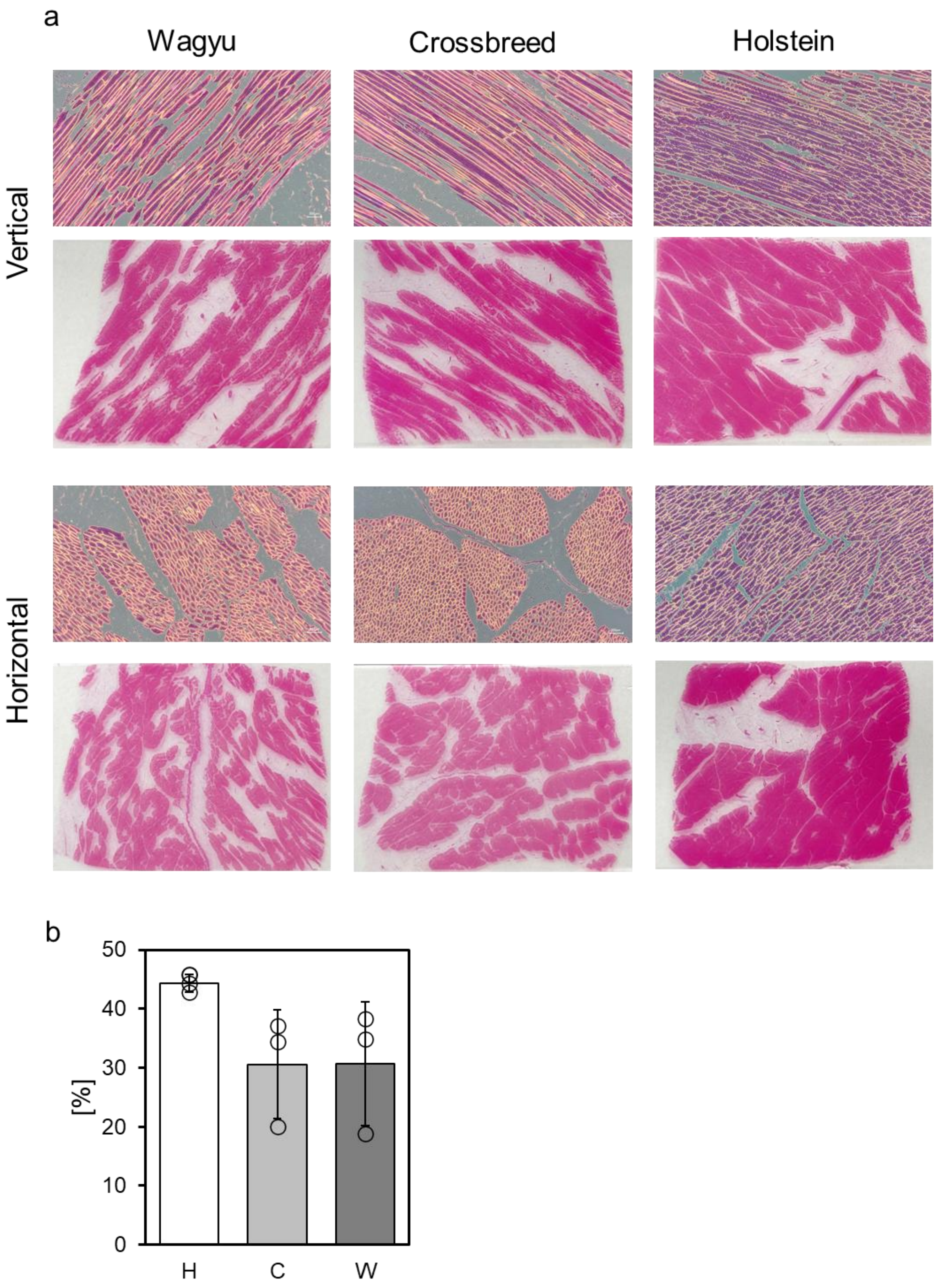


**Fig. S1. Histological analysis and weight loss rate by heating.**
**a,** Histological images by hematoxylin and eosin (HE) staining of Wagyu (W), Crossbreed (C) and Holstein (H) meats, respectively. **b,** Weight loss rate of each beef meet by heating (n=3).

a

**W vs C**

| Component | Flavor |
|---|---|
| Increased level in Wagyu | |
| 2,6-Dimethyl-5-heptenal | fruit, green, melon |
| Geraniol | rose, geranium |
| Decreased level in Wagyu | |
| 2,3-Pentanedione | cream, butter |

**W vs H**

| Component | Flavor |
|---|---|
| Increased level in Wagyu | |
| 3-Methylbutanal | malt |
| Butyl acetate | pear |
| Hexanal | grass, tallow, fat |
| 3-Pentanol | fruit |
| Pentyl acetate | banana |
| 2-Pentylfuran | green bean, butter |
| Hexyl acetate | fruit, herb |
| 2-Ethylpyridine | grass |
| 3-Heptanol | herb |
| trans-2-Heptenal | soap, fat, almond |
| Heptyl acetate | pear, fruity, aromatic, sweet |
| cis-3-Hexen-1-ol | grass |
| 2-Ethylhexyl acetate | fruit |
| Nonanal | fat, citrus, green |
| (Z)-hex-2-en-1-ol | leaf, green, wine, fruit |
| 1-Octen-3-ol | mushroom |
| 5-Ethyl-2,3-dimethylpyrazine | burnt, popcorn |
| Citronellal | fat |
| 2,4-Heptadienal | nut, fat |
| Benzaldehyde | almond, burnt sugar |
| 1-Octanol | chemical, metal, burnt |
| 2-Undecanone | orange, fresh, green |
| Undecanal | oil, pungent, sweet |
| (Z)-2-Decenal | tallow |
| gamma-Caprolactone | coumarin, sweet |
| n-Dodecanal | lily, fat, citrus |
| 2,6-Nonadienol | cucumber |
| 1-Undecanol | mandarin |
| 12-Methyltridecanal | cooked meat, tallow, fat, meat broth, sweat |
| (E)-Whiskey lactone | flower, lactone |
| gamma-Octalactone | coconut |
| Tetradecanal | flower, wax |
| Hydroxycitronellal | astringent, flowery, aldehyde |
| 1-Dodecanol | fat, wax |
| Phenol | phenol |
| gamma-Nonalactone | coconut, peach |
| Pentadecanal | fresh |
| Hexadecanal | cardboard |
| Massoia lactone | peach |
| Methyl palmitate | — |
| gamma-Undecalactone | apricot |
| Capric acid | fat |
| Farnesol isomer-2 | — |
| 1-Hexadecanol | wax, flower |
| Lauric acid | metal |
| Decreased level in Wagyu | |
| trans-3-Hexen-1-ol | moss, fresh |
| 2-Ethyl-5-methylpyrazine | fruit, sweet |

b

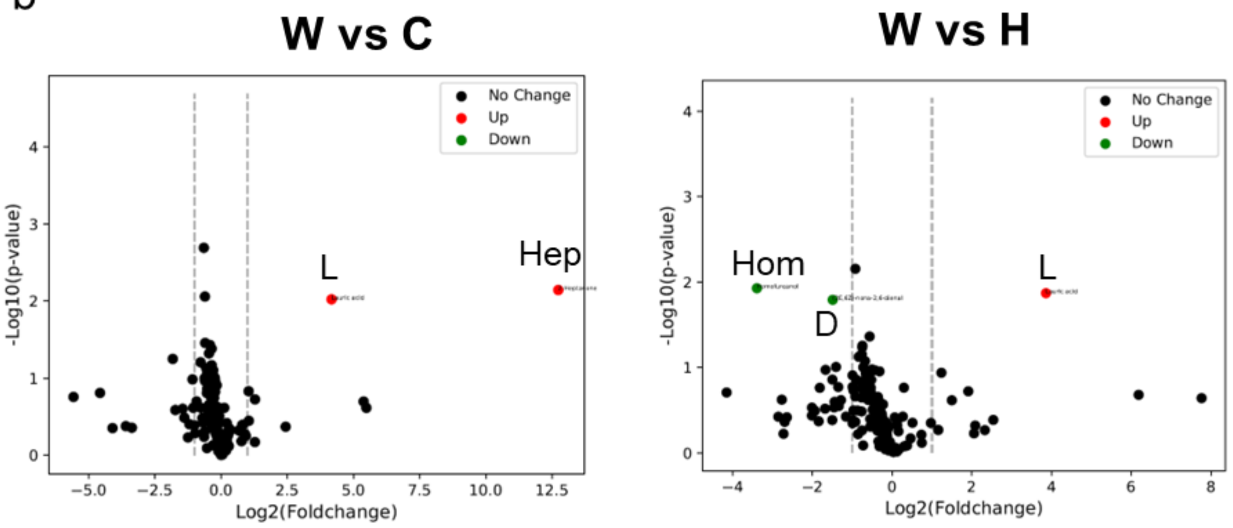


c

**W vs C**

| Component | Flavor |
|---|---|
| Increased level in Wagyu | |
| Lauric acid | metal |
| 2-Heptanone | soap |

**W vs H**

| Component | Flavor |
|---|---|
| Increased level in Wagyu | |
| Lauric acid | metal |
| Decreased level in Wagyu | |
| Homofureanol | caramel |
| (2E,6Z)-nona-2,6-dienal | cucumber, wax, green |

**Fig. S2. Summary of secreted aroma components from grilled meats.**

**a**, Summary of secreted aroma components from grilled fat and lean tissues. **b**, Volcano plots of secreted aroma components of grilled fat tissues in between W vs. C (left) and W vs. H (right), respectively. Hep: 2-Heptanone Lauric acid, Hom: Homofureanol, D:(2E,6Z)-nona-2,6-dienal, L: Lauric acid, respectively. **c**, Summary of secreted aroma components of grilled fat tissues in between W vs. C and W vs. H, respectively.

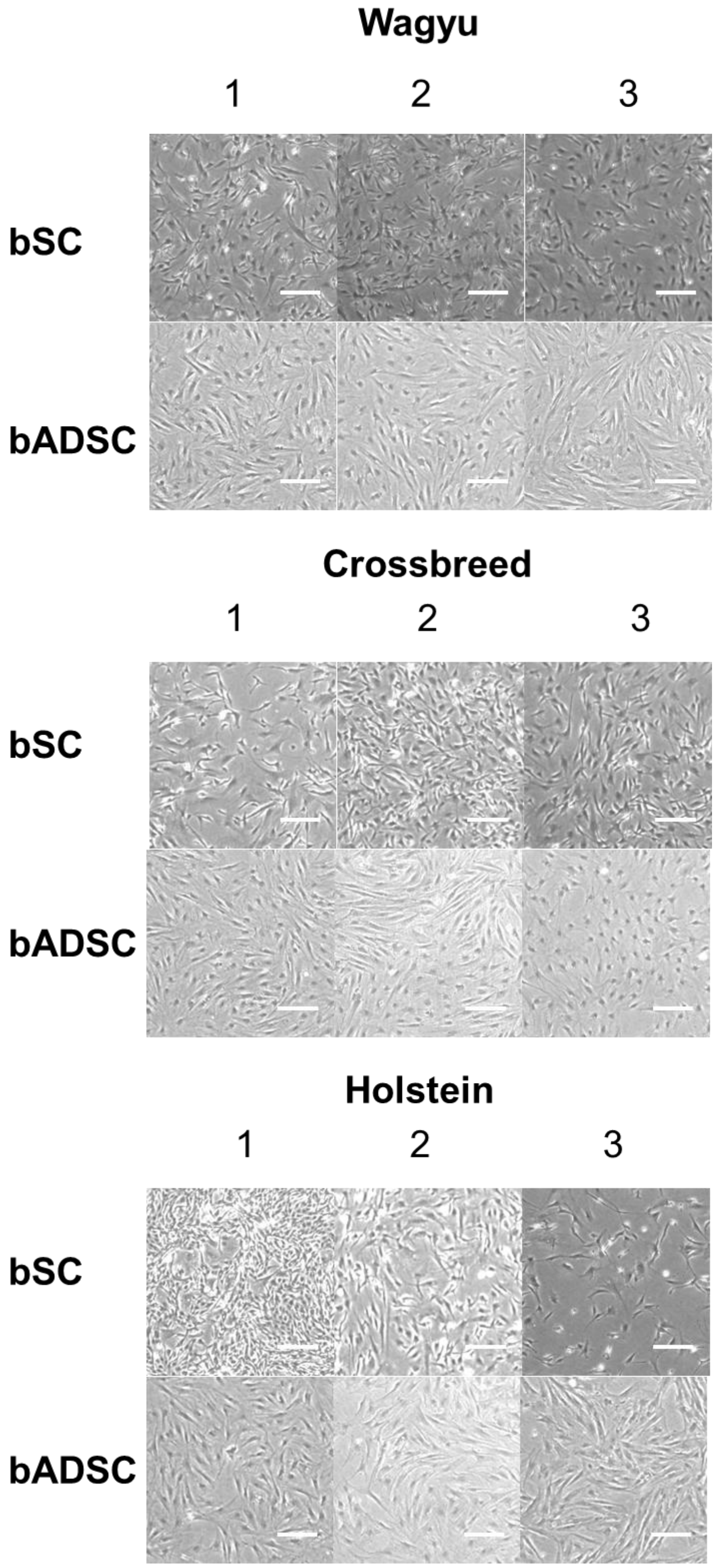


**Fig. S3. Phase contract image of bSC and bADSC.**

Phase contrast images of bSC and bADSC isolated from each cattle beef for fiber construction after 4 days of cultures (bSC) in DMEM 20%FBS and 7 days of cultures (bADSC) in DMEM 10%FBS, respectively. Scale bars are 200 μm.

## C vs W

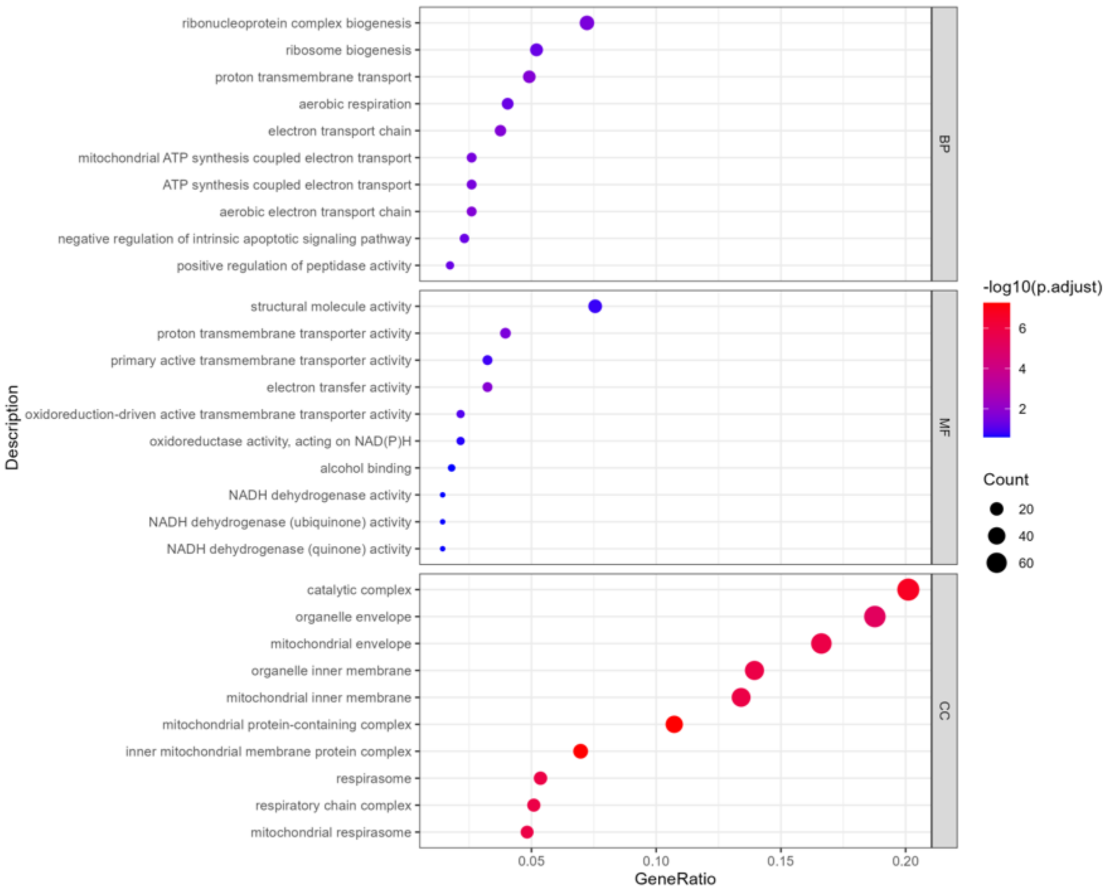

ribonucleoprotein complex biogenesis
ribosome biogenesis
proton transmembrane transport
aerobic respiration
electron transport chain
mitochondrial ATP synthesis coupled electron transport
ATP synthesis coupled electron transport
aerobic electron transport chain
negative regulation of intrinsic apoptotic signaling pathway
positive regulation of peptidase activity
structural molecule activity
proton transmembrane transporter activity
primary active transmembrane transporter activity
electron transfer activity
oxidoreduction-driven active transmembrane transporter activity
oxidoreductase activity, acting on NAD(P)H
alcohol binding
NADH dehydrogenase activity
NADH dehydrogenase (ubiquinone) activity
NADH dehydrogenase (quinone) activity
catalytic complex
organelle envelope
mitochondrial envelope
organelle inner membrane
mitochondrial inner membrane
mitochondrial protein-containing complex
inner mitochondrial membrane protein complex
respirasome
respiratory chain complex
mitochondrial respirasome
Description
BP
MF
CC
0.05
0.10
0.15
0.20
GeneRatio
-log10(p.adjust)
6
4
2
Count
20
40
60


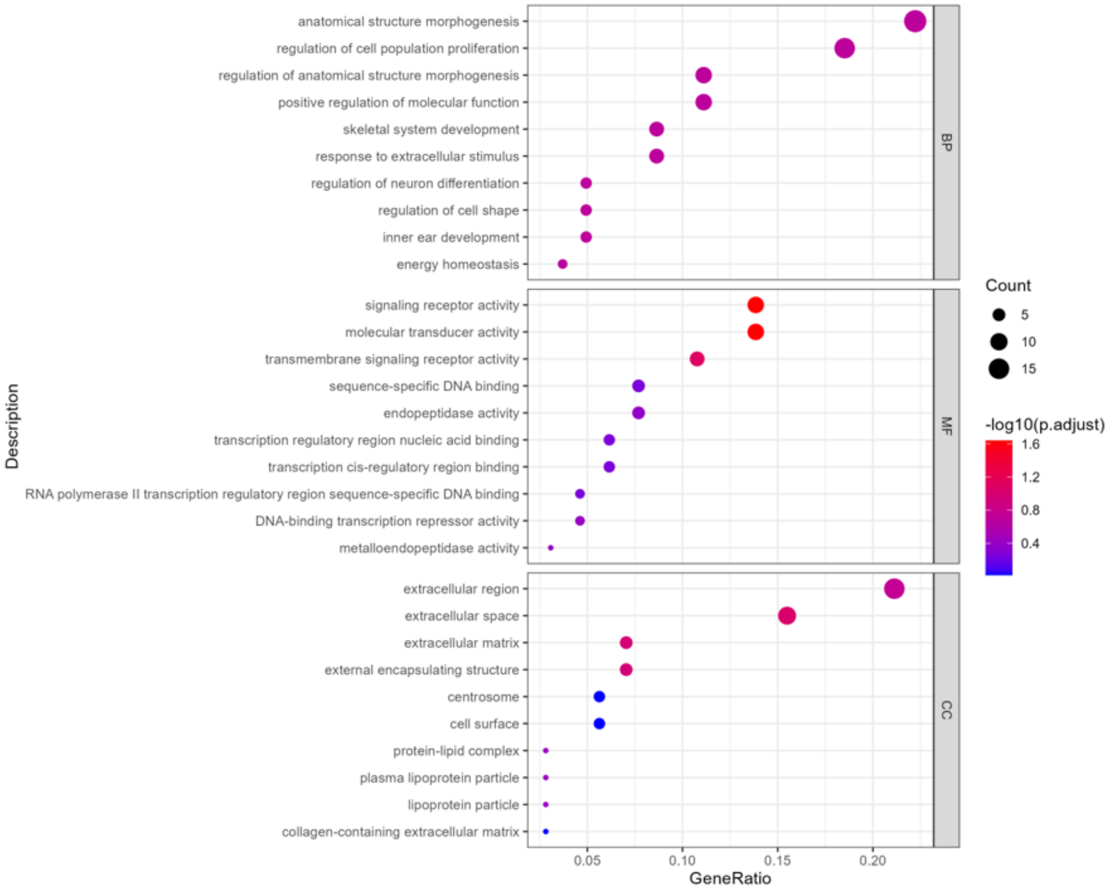

anatomical structure morphogenesis
regulation of cell population proliferation
regulation of anatomical structure morphogenesis
positive regulation of molecular function
skeletal system development
response to extracellular stimulus
regulation of neuron differentiation
regulation of cell shape
inner ear development
energy homeostasis
signaling receptor activity
molecular transducer activity
transmembrane signaling receptor activity
sequence-specific DNA binding
endopeptidase activity
transcription regulatory region nucleic acid binding
transcription cis-regulatory region binding
RNA polymerase II transcription regulatory region sequence-specific DNA binding
DNA-binding transcription repressor activity
metalloendopeptidase activity
extracellular region
extracellular space
extracellular matrix
external encapsulating structure
centrosome
cell surface
protein-lipid complex
plasma lipoprotein particle
lipoprotein particle
collagen-containing extracellular matrix
Description
BP
MF
CC
0.05
0.10
0.15
0.20
GeneRatio
Count
5
10
15
-log10(p.adjust)
1.6
1.2
0.8
0.4

**H vs W**

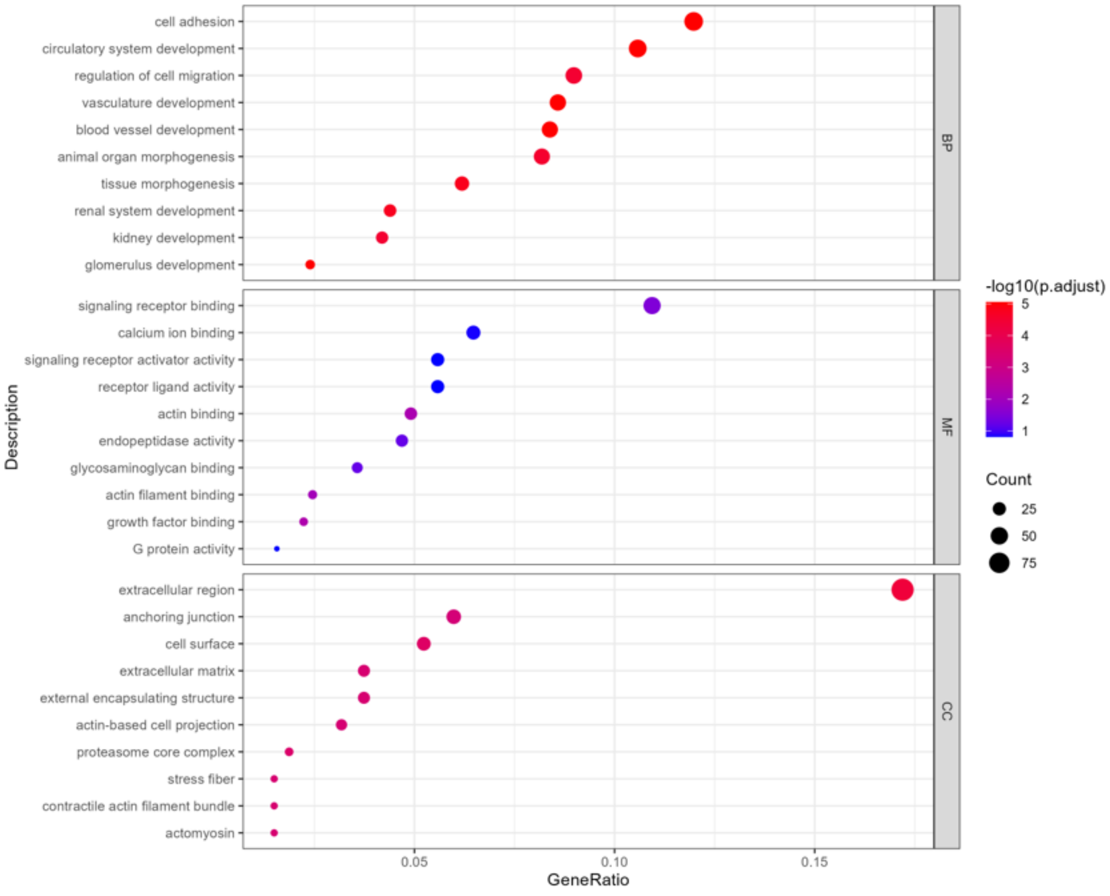
cell adhesion
circulatory system development
regulation of cell migration
vasculature development
blood vessel development
animal organ morphogenesis
tissue morphogenesis
renal system development
kidney development
glomerulus development
signaling receptor binding
calcium ion binding
signaling receptor activator activity
receptor ligand activity
actin binding
endopeptidase activity
glycosaminoglycan binding
actin filament binding
growth factor binding
G protein activity
extracellular region
anchoring junction
cell surface
extracellular matrix
external encapsulating structure
actin-based cell projection
proteasome core complex
stress fiber
contractile actin filament bundle
actomyosin
Description
BP
MF
CC
0.05
0.10
0.15
GeneRatio
-log10(p.adjust)
5
4
3
2
1
Count
25
50
75

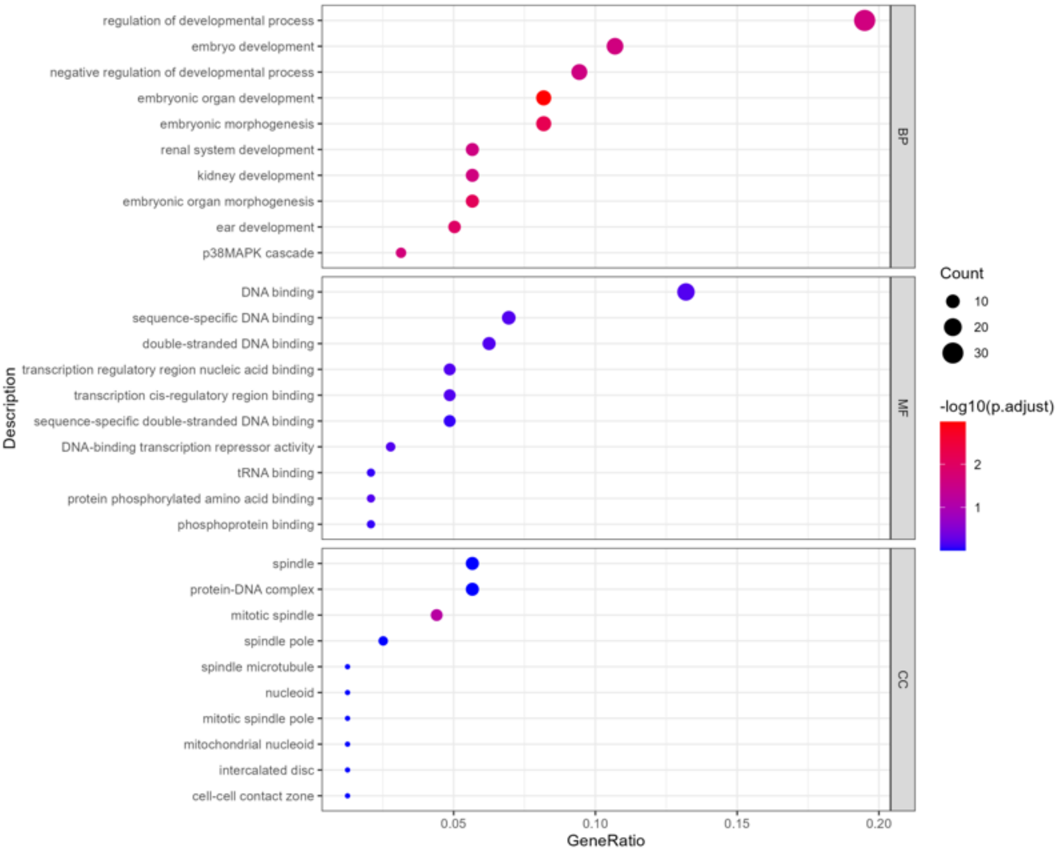
regulation of developmental process
embryo development
negative regulation of developmental process
embryonic organ development
embryonic morphogenesis
renal system development
kidney development
embryonic organ morphogenesis
ear development
p38MAPK cascade
DNA binding
sequence-specific DNA binding
double-stranded DNA binding
transcription regulatory region nucleic acid binding
transcription cis-regulatory region binding
sequence-specific double-stranded DNA binding
DNA-binding transcription repressor activity
tRNA binding
protein phosphorylated amino acid binding
phosphoprotein binding
spindle
protein-DNA complex
mitotic spindle
spindle pole
spindle microtubule
nucleoid
mitotic spindle pole
mitochondrial nucleoid
intercalated disc
cell-cell contact zone
Description
BP
MF
CC
0.05
0.10
0.15
0.20
GeneRatio
Count
10
20
30
-log10(p.adjust)
2
1

**Fig. S4. RNA sequence analyses of muscle and fat fibers.**

GO enrichment analysis of downregulated genes in cultivated muscle and fat fibers in the Crossbreed vs. Wagyu and Holstein vs. Wagyu comparisons.

**Table S1.** Comparison of RNA expression of fatty acid desaturase 1 (FADS1) and 2 (FADS2) from ADSCs and fat fibers from each cattle beef.

| FADS1 | sample_1 | sample_2 | value_1 | value_2 | log2(fold_change) | test_stat | p_value | q_value | significant |
|---|---|---|---|---|---|---|---|---|---|
| Crossbred_cell_ADSC_vs_Crossbre | Crossbred_cell_ADSC | Crossbred_fiber_ADSC | 7.05625 | 11.3571 | 0.686615 | 1.51758 | 0.00995 | 0.032552 | yes |
| Holstein_cell_ADSC_vs_Holstein_ | Holstein_cell_ADSC | Holstein_fiber_ADSC | 5.9085 | 9.88242 | 0.742071 | 1.80836 | 0.0019 | 0.009569 | yes |
| Wagyu_cell_ADSC_vs_Wagyu_fiber_ | Wagyu_cell_ADSC | Wagyu_fiber_ADSC | 7.12371 | 9.8825 | 0.472247 | 1.40397 | 0.0153 | 0.042177 | yes |
| FADS2 | | | | | | | | | |
| Crossbred_cell_ADSC_vs_Crossbre | Crossbred_cell_ADSC | Crossbred_fiber_ADSC | 0.543637 | 8.4003 | 3.94973 | 2.75148 | 0.00615 | 0.021861 | yes |
| Holstein_cell_ADSC_vs_Holstein_ | Holstein_cell_ADSC | Holstein_fiber_ADSC | 0.118974 | 5.13677 | 5.43215 | 2.17608 | 0.0044 | 0.018997 | yes |
| Wagyu_cell_ADSC_vs_Wagyu_fiber_ | Wagyu_cell_ADSC | Wagyu_fiber_ADSC | 0.303853 | 3.56115 | 3.5509 | 2.44159 | 0.00045 | 0.002001 | yes |

**Table S2.** Comparison of RNA expression of UCP1 and PRDM16 from fat fibers from each cattle beef.

| UCP1 | sample_1 | sample_2 | value_1 | value_2 | log2(fold_change) | test_stat | p_value | q_value | significant | Inverse of log2 |
|---|---|---|---|---|---|---|---|---|---|---|
| Crossbred_fiber_ADSC_vs_Wagyu_fiber_ADSC | Crossbred_fiber_ADSC | Wagyu_fiber_ADSC | 0.017685 | 0.027091 | 0.615262 | 0 | 1 | 1 | no | 1.625323846 |
| Holstein_fiber_ADSC_vs_Crossbred_fiber_ADSC | Holstein_fiber_ADSC | Crossbred_fiber_ADSC | 0 | 0.018101 | inf | 0 | 1 | 1 | no | - |
| Holstein_fiber_ADSC_vs_Wagyu_fiber_ADSC | Holstein_fiber_ADSC | Wagyu_fiber_ADSC | 0 | 0.027351 | inf | 0 | 1 | 1 | no | - |
| PRDM16 | | | | | | | | | | |
| Crossbred_fiber_ADSC_vs_Wagyu_fiber_ADSC | Crossbred_fiber_ADSC | Wagyu_fiber_ADSC | 0.479807 | 0.693195 | 0.530807 | 1.16857 | 0.03395 | 0.242668 | no | 1.883923912 |
| Holstein_fiber_ADSC_vs_Crossbred_fiber_ADSC | Holstein_fiber_ADSC | Crossbred_fiber_ADSC | 0.350145 | 0.48981 | 0.48427 | 0.892992 | 0.1007 | 0.794157 | no | 2.06496376 |
| Holstein_fiber_ADSC_vs_Wagyu_fiber_ADSC | Holstein_fiber_ADSC | Wagyu_fiber_ADSC | 0.350044 | 0.701686 | 1.00329 | 1.89989 | 0.00115 | 0.026368 | yes | 0.996720789 |

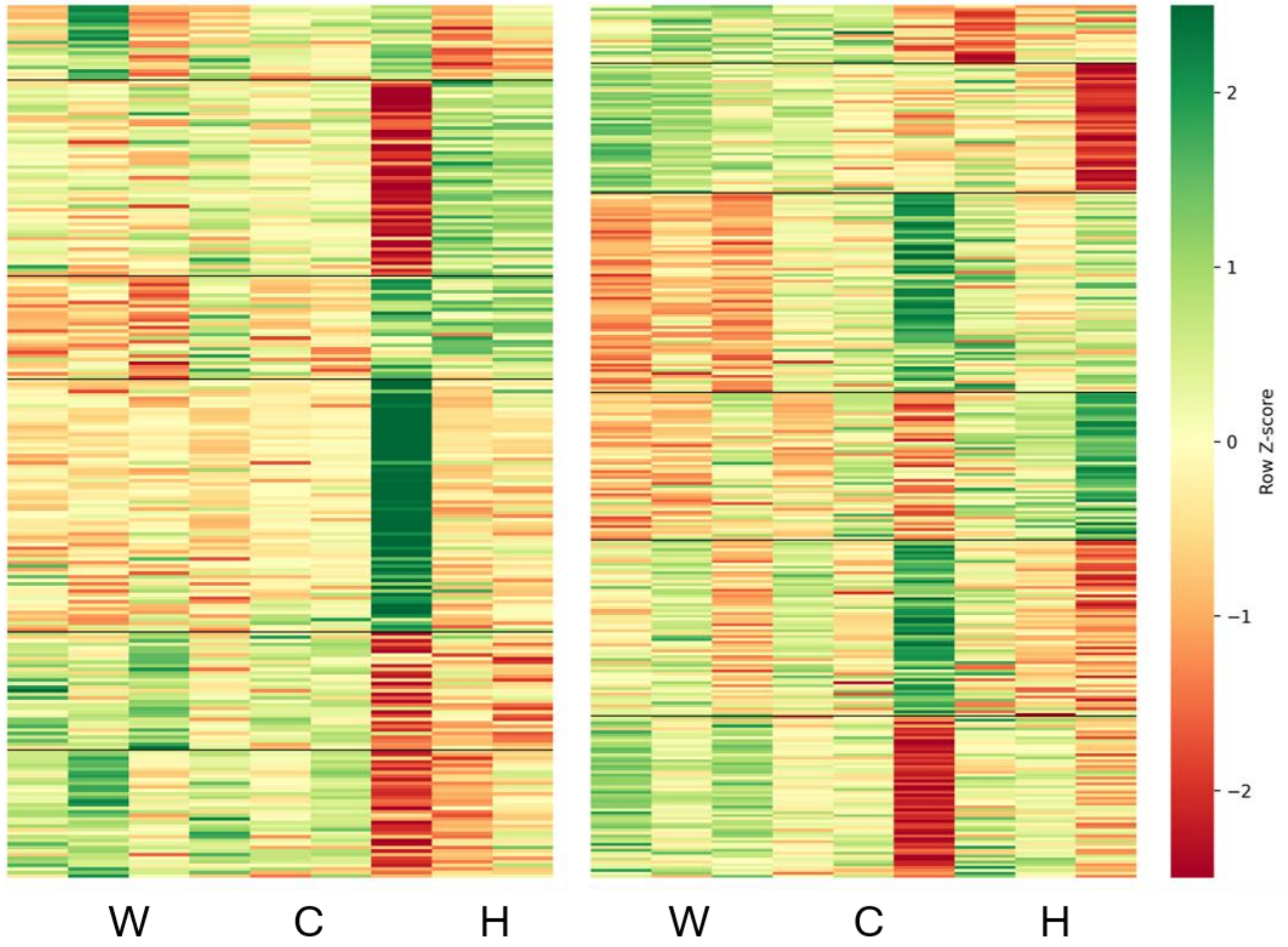


**Fig. S5. Clustering of gene expression profiles using GO and Hallmark gene sets in bSC and bADSC.**

Heatmaps show expression entries in bSC (left, k = 6, 245 entries) and bADSC (right, k = 6, 302 entries). Candidate genes were selected from GO Biological Process and Hallmark gene sets related to myogenesis, adipocyte differentiation, and fatty acid metabolism. Entries passing the expression and variance filters were $\log_2$(FPKM+1)-transformed, standardized using row-wise Z-scores across nine samples, and independently partitioned into six clusters for each cell type using k-means. Rows represent individual expression entries, with entries sharing the same gene symbol retained separately. Black horizontal lines indicate cluster boundaries. Green and red indicate relatively higher and lower expression within each row, respectively. *k* denotes the number of clusters, and *n* denotes the total number of expression entries included in the analysis.

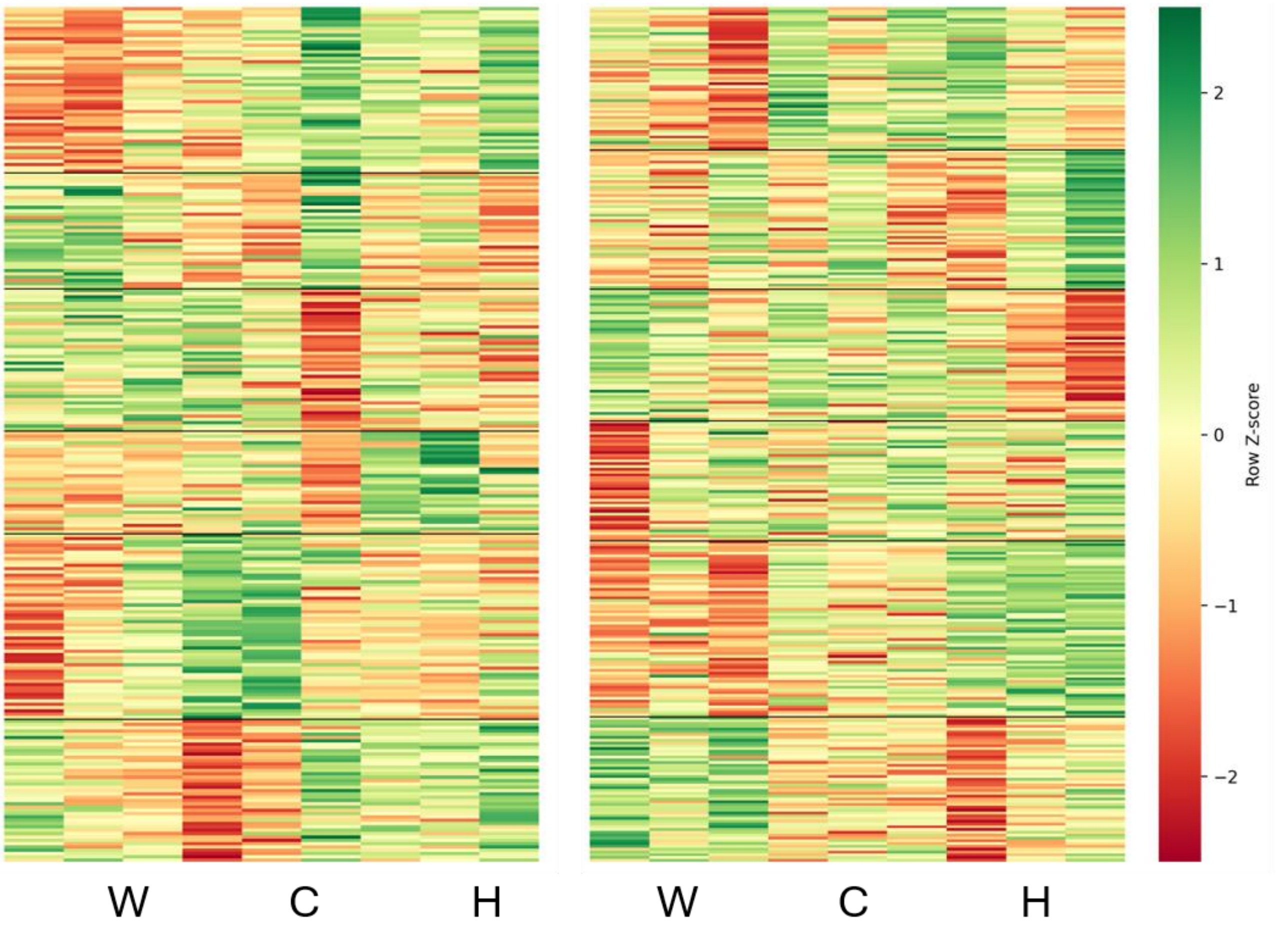


**Fig. S6. Clustering of gene expression profiles using GO and Hallmark gene sets in muscle and fat fibers.**

Heatmaps show expression entries in muscle fibers (left, k = 6, 258 entries) and fat fibers (right, k = 6, 306 entries). Candidate genes were selected from GO Biological Process and Hallmark gene sets related to myogenesis, adipocyte differentiation, and fatty acid metabolism. Entries passing the expression and variance filters were $\log_2$(FPKM+1)-transformed, standardized using row-wise Z-scores across nine samples, and independently partitioned into six clusters for each cell type using k-means. Rows represent individual expression entries, with entries sharing the same gene symbol retained separately. Black horizontal lines indicate cluster boundaries. Green and red indicate relatively higher and lower expression within each row, respectively. *k* denotes the number of clusters, and *n* denotes the total number of expression entries included in the analysis.

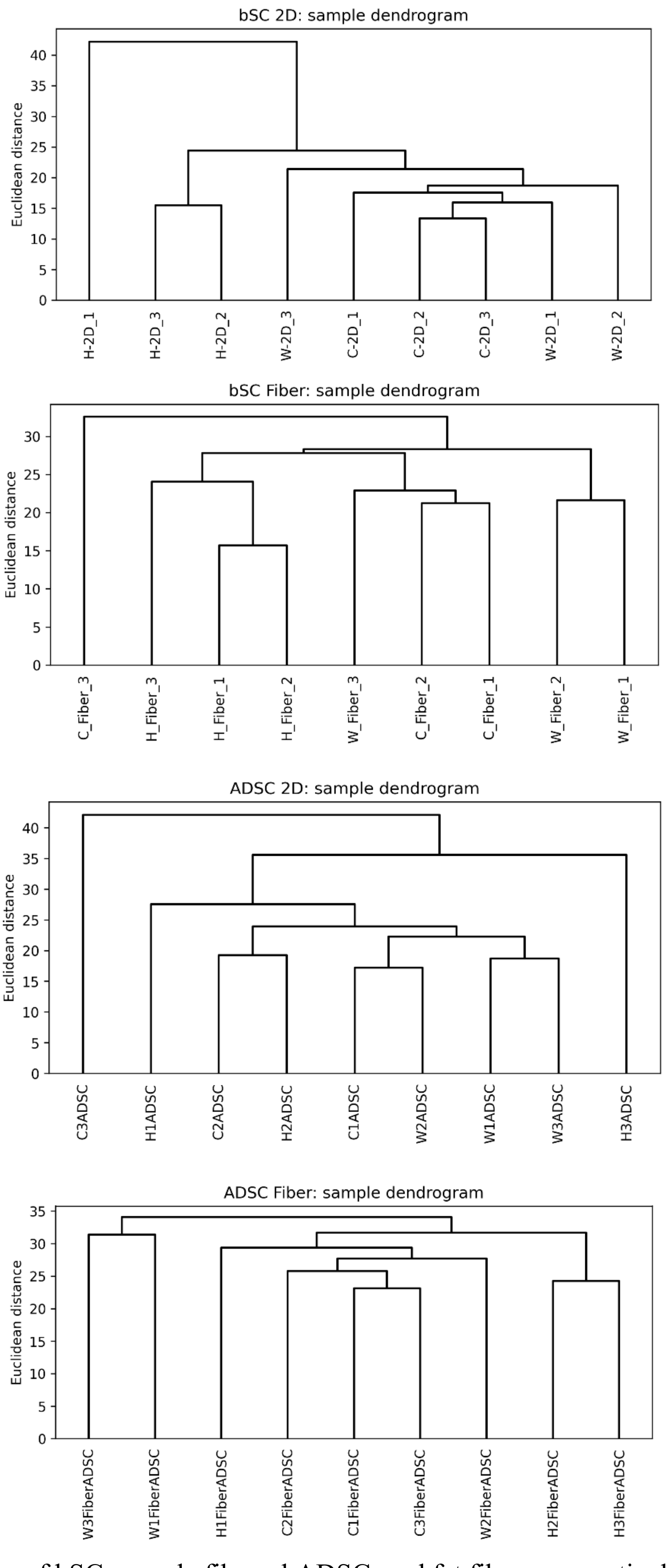


**Fig. S7.** Dendrograms of bSC, muscle fibers, bADSC, and fat fibers, respectively.

**Methods. Clustering of gene expression profiles using GO and Hallmark gene sets**

Clustering of gene expression profiles using GO and Hallmark gene sets was performed separately for bSCs and bADSCs. FPKM expression values from three replicates per condition (W, C, and H) were analyzed. For bSCs, only samples from the two-dimensional culture condition were included; Fiber samples were excluded from both analyses.

Candidate genes were obtained from the human GO Biological Process and Hallmark collections of the Molecular Signatures Database (MSigDB). The selected gene sets were HALLMARK_MYOGENESIS, GOBP_SKELETAL_MUSCLE_CELL_DIFFERENTIATION, GOBP_MYOBLAST_DIFFERENTIATION, and GOBP_MUSCLE_CONTRACTION for bSCs, and HALLMARK_ADIPOGENESIS, GOBP_FAT_CELL_DIFFERENTIATION, and GOBP_FATTY_ACID_METABOLIC_PROCESS for ADSCs. The union of genes within the selected sets was generated separately for each cell type.

Human candidate genes were mapped to bovine genes using orthology information retrieved from Ensembl BioMart (Ensembl Genes release 116; bovine reference assembly ARS-UCD2.0). Only one-to-one orthologs with non-empty human and bovine gene symbols were retained. Bovine gene symbols associated with multiple Ensembl gene identifiers were excluded to avoid ambiguous matching. Candidate bovine genes were matched to the FPKM expression tables by gene symbol.

Expression entries lacking gene symbols were excluded. Original XLOC identifiers were retained as unique row identifiers, and entries sharing the same gene symbol were analyzed separately without averaging or summation. Thus, clustering was performed at the level of XLOC-defined expression entries rather than aggregated gene symbols. Entries were retained if they exhibited FPKM $\geq 1$ in at least three of the nine samples. Expression values were transformed as $\log_2(\mathrm{FPKM} + 1)$, and entries with a variance of $\leq 10^{-12}$ across samples were excluded. When more than 1,000 entries passed these filters, the 1,000 entries with the highest variance in log-transformed expression were selected; otherwise, all eligible entries were used.

The log-transformed expression values were standardized independently for each entry across the nine samples using row-wise Z-scores, with the standard deviation calculated using a divisor of n (ddof = 0). K-means clustering was performed using scikit-learn with six clusters, 30 initializations, and a fixed random seed of 42. Inertia and silhouette scores were additionally calculated for candidate cluster numbers ranging from 2 to 10 as diagnostic measures.

Heatmaps displayed row-wise Z-scores for individual samples, with entries grouped by cluster assignment. For cluster-profile plots, row-wise Z-scores were displayed for all nine individual samples. Individual entry profiles and cluster mean profiles were shown; replicate values were not averaged within conditions. Clustering was performed using all nine individual samples. Sample dendrograms were generated independently for each cell type (bSCs and ADSCs) and culture condition (2D and Fiber) using the row-wise Z-scores of $\log_2(\mathrm{FPKM} + 1)$ values for expression entries selected independently within each analysis as described above. Analyses and visualizations were conducted in Python 3.11.8 using pandas 2.2.1, NumPy 1.26.4, scikit-learn 1.4.1.post1, Matplotlib 3.8.3, and seaborn 0.12.2.
